# Non-canonical Wnt signaling triggered by WNT2B drives adrenal aldosterone production

**DOI:** 10.1101/2024.08.23.609423

**Authors:** Kleiton S. Borges, Donald W. Little, Taciani de Almeida Magalhães, Claudio Ribeiro, Typhanie Dumontet, Chris Lapensee, Kaitlin J. Basham, Aishwarya Seth, Svetlana Azova, Nick A. Guagliardo, Paula Q. Barrett, Mesut Berber, Amy E. O’Connell, Adina F. Turcu, Antonio Marcondes Lerario, Dipika R. Mohan, William Rainey, Diana L. Carlone, Joel N. Hirschhorn, Adrian Salic, David T. Breault, Gary D. Hammer

## Abstract

The steroid hormone aldosterone, produced by the zona glomerulosa (zG) of the adrenal gland, is a master regulator of plasma electrolytes and blood pressure. While aldosterone control by the renin-angiotensin system is well understood, other key regulatory factors have remained elusive. Here, we replicated a prior association between a non-coding variant in *WNT2B* and an increased risk of primary aldosteronism, a prevalent and debilitating disease caused by excessive aldosterone production. We further show that in both mice and humans, WNT2B is expressed in the mesenchymal capsule surrounding the adrenal cortex, in close proximity to the zG. Global loss of *Wnt2b* in the mouse results in a dysmorphic and hypocellular zG, with impaired aldosterone production. Similarly, humans harboring *WNT2B* loss-of-function mutations develop a novel form of Familial Hyperreninemic Hypoaldosteronism, designated here as Type 4. Additionally, we demonstrate that WNT2B signals by activating the non-canonical Wnt/planar cell polarity pathway. Our findings identify WNT2B as a key regulator of zG function and aldosterone production with important clinical implications.

**Highlights:**

- *WNT2B* variant is associated with increased risk for primary aldosteronism
- *Wnt2b* knock-out mice show defects in adrenal morphology
- *Wnt2b* knock-out mice have hyperreninemic hypoaldosteronism
- WNT2B activates non-canonical Wnt/planar cell polarity signaling
- WNT2B deficiency causes a new form of familial hyperreninemic hypoaldosteronism

## Introduction

The adrenal gland is encapsulated by a mesenchymal cell layer and contains an underlying cortex, which is divided into three distinct zones: the outermost zona glomerulosa (zG), the zona fasciculata (zF), and the innermost zona reticularis (zR)(1). The cells of the zG, organized into rosettes(2–4), produce aldosterone, the mineralocorticoid hormone that regulates sodium homeostasis and concomitant intravascular volume, under the control of the renin-angiotensin-aldosterone system (RAAS) and extracellular potassium(1, 5). Dysfunction of the zG leads to distinct human diseases with outcomes determined by the resultant levels of aldosterone production(6–11). Insufficient aldosterone production leads to hypoaldosteronism (Hypo-A), which can be familial (caused by CYP11B2 loss-of-function (LOF) leading to Aldosterone Synthase deficiency or LGR4 LOF leading to defects in the R-spondin 4 receptor)(8, 9) or acquired (such as caused by autoimmunity, infections or medications that target the RAAS)(12). Conversely, autonomous aldosterone production by the zG results in primary aldosteronism (PA)(11, 13). PA is the most common form of endocrine hypertension, affecting 8-10% of patients with hypertension, and it is associated with a higher risk of cardiovascular and renal damage compared to primary hypertension of similar severity(11, 13). While genetic studies have identified somatic mutations in various ion transport genes that result in depolarization-driven increases in aldosterone production(11, 14), these genetic alterations do not account for the zG hyperplasia observed in some patients with PA(15–19).

Wnt signaling is triggered by the WNT family of secreted ligands, and orchestrates numerous developmental processes, including cell fate determination, differentiation, proliferation and migration(20–22). While all WNTs signal through Frizzled receptors (FZDs), they are divided into canonical and non-canonical WNTs, depending on the coreceptors they engage(23). Canonical WNTs use low-density lipoprotein coreceptors, LRP5 and LRP6, and signal through β-catenin to control target gene expression. Non-canonical WNTs use the tyrosine-protein kinase coreceptors, including ROR1 and ROR2, to activate Ca^2+^ signaling or signal through small GTPases to control planar cell polarity (PCP)(24–30).

Wnt signaling plays crucial roles in zG development and function(3, 31–40). For instance, mesenchymal capsular cells express R-spondin 3 (RSPO3), a ligand for LGR receptors, which is a strong potentiator of canonical Wnt/β-catenin signaling in the zG(41). Increasing or reducing β-catenin levels has strong effects on zG morphology, pointing to the importance of canonical Wnt signaling in zG homeostasis(3, 8, 31, 34–41). Additionally, gain-of-function (GOF) mutations in *CTNNB1*, which encodes β-catenin, act as driver mutations in PA(10, 42–45). Finally, recent genome-wide association (GWAS) studies identified a non-coding common variant in *WNT2B* (rs3790604) that correlates with the highest risk of developing PA(46, 47). Despite this evidence, the cellular and molecular mechanisms by which WNT2B controls adrenal function remain unknown.

Here, we demonstrate an essential role for WNT2B and non-canonical Wnt/PCP signaling in zG function and aldosterone production. We validate the association of *WNT2B* variant rs3790604 with increased risk of developing PA using an independent case-controlled multi-ancestry cohort from the *All of Us* Database(48, 49). We show that mice lacking WNT2B have impaired aldosterone production, compensated by elevated levels of plasma renin, establishing WNT2B as a key regulator of zG function. Additionally, we show that WNT2B activates non-canonical Wnt signaling. In both human and mouse adrenals, WNT2B is expressed in the adrenal capsule, while the non-canonical Wnt pathway components are enriched in the zG, suggesting that WNT2B signals to the underlying zG. Finally, we show that humans with LOF mutations in *WNT2B* exhibit a novel form of Familial Hyperreninemic Hypoaldosteronism, designated here as Type 4. Our findings identify WNT2B as a key activator of zG function and aldosterone production linked to human adrenal disease.

## Results

### Non-coding Variant in *WNT2B* is associated with increased risk of PA

Recent multi-ancestry GWAS meta-analyses identified a common non-coding variant in *WNT2B* (rs3790604) that is associated with risk of developing PA(46, 47). To replicate this finding, we performed an independent case-controlled multi-ancestry cohort GWAS using the *All of Us* Database(48, 49). Among the 245,195 participants in the database with short read whole genome sequence data, we identified 271 cases of PA and matched them by genetic ancestry to five controls each, drawn from 74,354 controls with no documented record of hypertension or elevated blood pressure. We then tested the association between the rs3790604 variant and PA using logistic regression models. Despite the low number of cases, we nominally replicated the association of the A allele of rs3790604 with increased risk of PA (odds ratio of 1.53, 95% confidence interval 1.06-2.20, one-tailed p-value=0.01). The observed odds ratio is consistent with prior observations, strengthening the previous conclusion that carriage of the risk allele correlates with the development of PA. Because it is not known how WNT2B might influence the development of PA, we investigated the connection between WNT2B and aldosterone production in mice and humans.

### *Wnt2b* is required for zG formation and maintenance

To explore the role of WNT2B in adrenocortical function, we first employed single molecule *in situ* hybridization (RNAscope) to assess *Wnt2b* expression in the adult mouse adrenal. *Wnt2b* transcripts were exclusively found in the mesenchymal capsule (Fig. 1a), consistent with its expression pattern during adrenal development(50, 51). To investigate the functional role of WNT2B in the adrenal, we generated whole body knock-out mice (*Wnt2b^-/-^*)(52) (Fig. 1b, Supplemental Figure 1a). In both female and male mice, *Wnt2b* deletion resulted in an ∼25% reduction in adrenal weight compared to wild type (WT) adrenals (Fig. 1c, Supplemental Figure 1b) and a marked disruption of rosette structures in the outer adrenal cortex (dotted white lines), a hallmark of zG morphology(2–4)(Fig. 1d, Supplemental Figure 1c). Indeed, using immunofluorescence staining for LAMB1, which delineates rosette boundaries in the adult zG(3), rosette structures were essentially absent from *Wnt2b^-/-^* adrenals compared to WT adrenals (Fig. 1e). Moreover, immunofluorescent staining showed a marked reduction in the number of cells expressing the zG-specific markers DAB2(53) and aldosterone synthase (CYP11B2)(5), and a near complete loss of Gαq(34) and β-catenin(36) in *Wnt2b^-/-^*compared to WT adrenals, suggesting a near complete lack of the zG layer (Fig. 1f, Supplemental Figure 1d). To better delineate differences in zonation of the adrenal cortex we performed co-staining of *Wnt2b^-/-^* and WT adrenals for DAB2 and the zF-specific marker AKR1B7(54). In select regions of *Wnt2b^-/-^* adrenals, AKR1B7-positive/DAB2-negative cells extended to the adrenal capsule (Supplemental Figure. 1e), further confirming the marked reduction in the number of zG cells in *Wnt2b^-/-^*mice. We next performed gene expression analysis of *Wnt2b^-/-^* adrenals, which confirmed reduced expression of zG-specific genes, such as *Cyp11b2* and *Dab*2, as well as the β-catenin target genes *Wnt4* and *Lef1*(37) (Supplemental Figure 1f-g). In addition, we observed a marked decrease in the expression of *Shh* (Supplemental Figure 1f-g), a zG-specific gene important for adrenocortical development and steroidogenic progenitor cells, which signals to the overlying capsule to regulate *Gli1* expression(33, 55–57). This decrease was accompanied by a reduced thickness of the capsule in Wnt2b^-/-^ adrenals, confirmed by immunostaining for the capsule-specific marker NR2F2 (COUP-TFII)(41, 58) (Supplemental Figure 1h) and downregulation of *Gli1* expression (Supplemental Figure 1f-g). Taken together, these data indicate that WNT2B plays a critical role in zG formation and is important for signaling in the adrenal.

**Figure 1.**
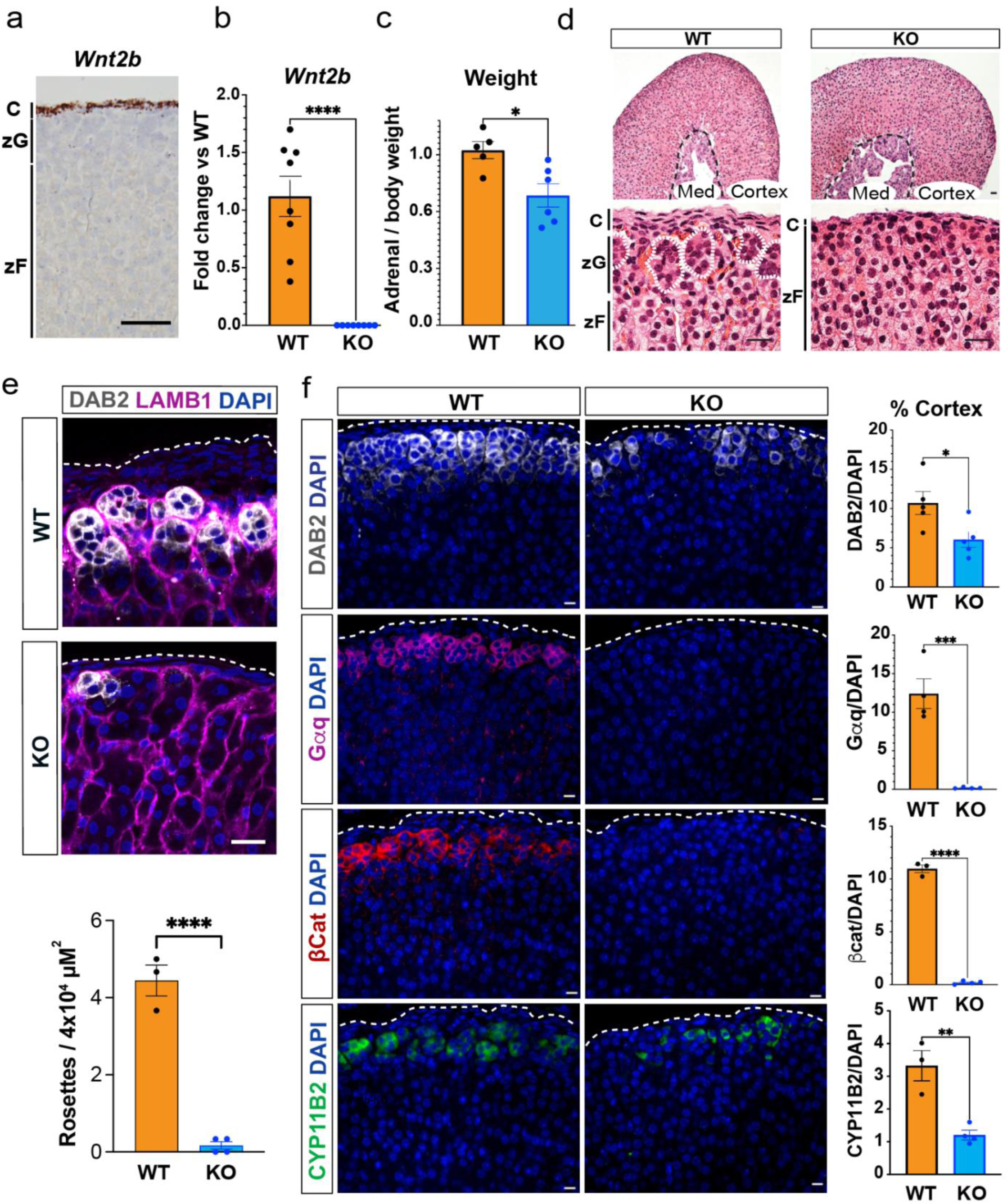
WNT2B deficiency results in a dysmorphic zG in mice. a. Representative image of *Wnt2b* expression using single molecule *in situ* hybridization (smISH) from an adult female adrenal. Scale bar: 100μm. C, capsule; zG, zona glomerulosa; zF, zona fasciculata. b. QRT-PCR was performed on WT and KO female adrenals (n=8 WT, n=8 KO). Two-tailed Student’s t-test. ****p < 0.0001. Data are represented as mean ± SEM. c. Adrenal weight normalized to body weight from female mice (n=5 WT, n=6 KO). Two-tailed Student’s t-test. *p < 0.05. Data are represented as mean fold change ± SEM. d. Representative H&E images of WT and KO female adrenals. Scale bar: 10μm. Dotted white line delineates rosette structures. C, capsule; zG, zona glomerulosa; zF, zona fasciculata. Med, medulla. e. Representative image and quantification of immunohistochemistry from WT and KO female adrenals stained for Laminin β1 (LAMB1, magenta), indicating the basement membrane surrounding distinct clusters of zG cells (DAB2, gray; DAPI, blue). Scale bar: 20μm. Staining delineates individual glomeruli, highlighting the loss of rosettes in KO adrenals. The number of DAB2+ clusters containing ≥5 cells (n=3 WT, n=4 KO). Two-tailed Student’s t-test. ****p < 0.0001. Data are represented as mean ± SEM. f. Representative images and quantification from female adrenals immunostained for DAB2 (gray, n=5 WT, n=5 KO), Gαq (magenta, n=4 WT, n=4 KO), β-catenin (β-cat, red, n=3 WT, n=4 KO) and CYP11B2 (green, n=3 WT, n=4 KO). Positive cells were quantified and normalized to nuclei (DAPI, blue) in the cortex. Scale bars: 10μm. Two-tailed Student’s t-test. *p < 0.05; **p < 0.01; ***p < 0.001; ****p < 0.0001. Data are represented as mean ± SEM.

To determine if WNT2B also functions in maintaining the zG in adult mice, we tested whether genetic ablation of *Wnt2b* specifically within the adrenal capsule similarly disrupts zG morphology. To delete *Wnt2b* in the adult, we generated conditional *Gli1^CreER/+^* :: *Wnt2b^fl/fl^* mice, which allowed for temporal control of Cre recombination in Gli1+ capsular cells(59) with tamoxifen. Adult mice were treated with tamoxifen at six weeks of age and adrenal glands were assessed four weeks later (Supplemental Figure 1i). As expected, *Wnt2b* conditional knock-out (cKO) mice exhibit a significant (∼90%) reduction in adrenal *Wnt2b* expression compared to control mice (Supplemental Figure 1j). Remarkably, expression of *Cyp11b2* was reduced by ∼40% in cKO adrenals compared to controls (Supplemental Figure 1k). To establish the effect of cKO on zG morphology, we performed immunostaining for CYP11B2 and DAB2, which revealed a marked reduction in both markers (Supplemental Figure 1l). Moreover, immunostaining for the zF-specific marker CYP11B1(60) revealed expression extended to the capsule, underscoring the decreased size of the zG in cKO mice (Supplemental Figure 1m). Together, these findings align with the marked reduction in the number of zG cells observed in global *Wnt2b^-/-^* mice and show that *Wnt2b* is also essential for zG maintenance in the adult.

### Loss of *Wnt2b* causes hypoaldosteronism

Because the zG is the source of circulating aldosterone, we evaluated plasma aldosterone levels in both *Wnt2b^-/-^*and WT mice. Despite a pronounced disruption in the zG layer and a notable reduction in the number of aldosterone-producing cells (Fig. 1f, Supplemental Figure 1d), we observed no significant difference in plasma aldosterone levels between the two groups (Fig. 2a). However, plasma renin concentration in *Wnt2b^-/-^* mice was significantly elevated, indicative of increased RAAS activation and thus a state of compensated hypoaldosteronism (Hypo-A)(7) (Fig. 2b). Importantly, levels of plasma corticosterone produced from the zF were not different between *Wnt2b^-/-^* and WT mice (Supplemental Figure 2), indicating that the observed phenotype was not the result of a global defect in adrenal steroid production. To better understand the mechanisms supporting aldosterone levels in *Wnt2b^-/-^* mice (Fig. 2a) despite reduced numbers of zG cells (Fig. 1f, Supplemental Figure 1d), we analyzed aldosterone and corticosterone secretion in WT and *Wnt2b^-/-^* adrenal slice cultures *ex vivo*(61). This analysis revealed a markedly decreased rate of aldosterone secretion, but an unchanged rate of corticosterone secretion from *Wnt2b^-/-^* adrenals, resulting in a decreased aldosterone/corticosterone ratio compared to WT (Fig. 2c). These findings confirm an autonomous defect in aldosterone production in *Wnt2b^-/-^* adrenals, consistent with the decreased number of zG cells. This defect leads to reduced aldosterone secretion, which *in vivo* is counterbalanced by a compensatory increase in plasma renin levels.

**Figure 2.**
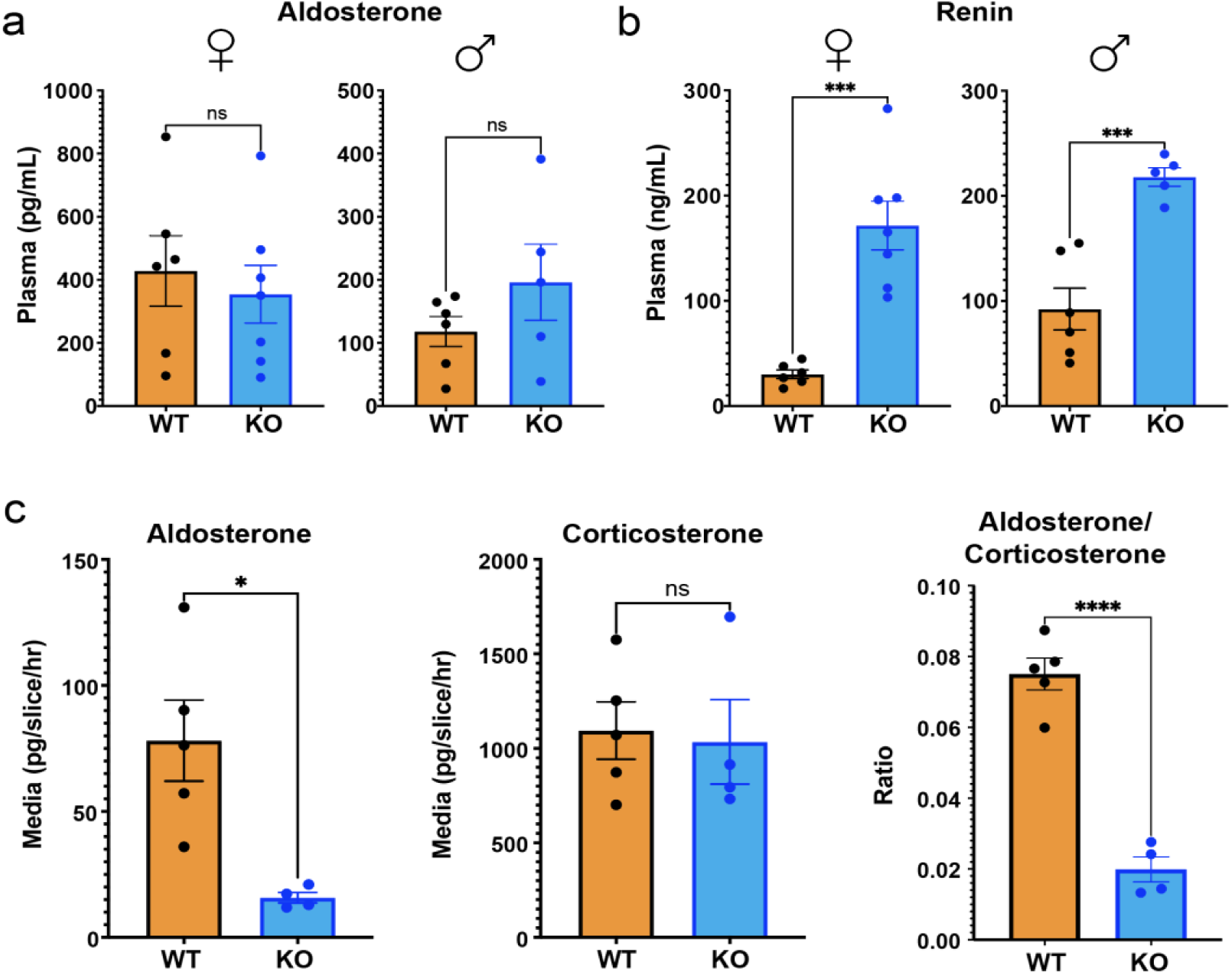
WNT2B deficiency results in hypoaldosteronism in mice. a. Quantification of plasma aldosterone levels (female, n=6 WT, n=7 KO; male, n=6 WT, n=5 KO). Two-tailed Student’s t-test. ns, not significant. Data are represented as mean ± SEM. b. Quantification of plasma renin levels (female, n=6 WT, n=7 KO; male n=6 WT, n=5 KO). Two -tailed Student’s t-test. ***p < 0.001. Data are represented as mean ± SEM. c. Aldosterone, corticosterone and aldosterone/corticosterone ratios produced from male adrenal slice preparations, *ex vivo*, plotted as mean of each mouse (n=5 WT, n=4 KO). Two-tailed Student’s t-test. *p < 0.05, ****p < 0.0001, ns, not significant. Data are represented as mean ± SEM.

### Activation of canonical Wnt/β-catenin signaling fails to rescue WNT2B deficiency

The significant decrease in expression of canonical Wnt/β-catenin target genes in *Wnt2b^-/-^* mouse adrenals (Supplemental Figure 1f-g) suggested that WNT2B may function as a canonical WNT ligand. Thus, we tested whether the *Wnt2b^-/-^* adrenal phenotype could be rescued by activating canonical Wnt/β-catenin signaling through chronic administered of lithium chloride (LiCl), which activates canonical WNT signaling by inhibiting GSK3β, leading to β-catenin stabilization(62, 63). When mice were treated with LiCl from birth for 6 weeks (Fig. 3a), LEF-1 expression was extensively induced in the adrenal cortex of *Wnt2b^-/-^* mice compared to untreated mice, confirming activation of the canonical Wnt/β-catenin pathway (Supplemental Figure 3). Moreover, expression was high even in the zF where LEF-1 is not typically expressed(3) (Supplemental Figure 3), consistent with stabilization of pre-existing β-catenin. Notably, expression of β-catenin and DAB2 were induced in the outer adrenal cortex of LiCl-treated *Wnt2b^-/-^* mice (Fig. 3b). In contrast, Gαq and CYP11B2, both required for aldosterone production(5, 64), were not restored by LiCl treatment (Fig. 3b), indicating that zG morphology was not fully rescued. Consistent with this, LiCl-treated *Wnt2b^-/-^*mice exhibited the same plasma levels of aldosterone and renin observed in the untreated *Wnt2b^-/-^* mice, confirming that chronic activation of the canonical Wnt/β-catenin pathway was insufficient to rescue the functional defect in zG activity caused by Wnt2b loss (Fig. 3c-d), indicating a potential role for WNT2B in activating a non-canonical WNT pathway to govern zG morphogenesis and function.

**Figure 3.**
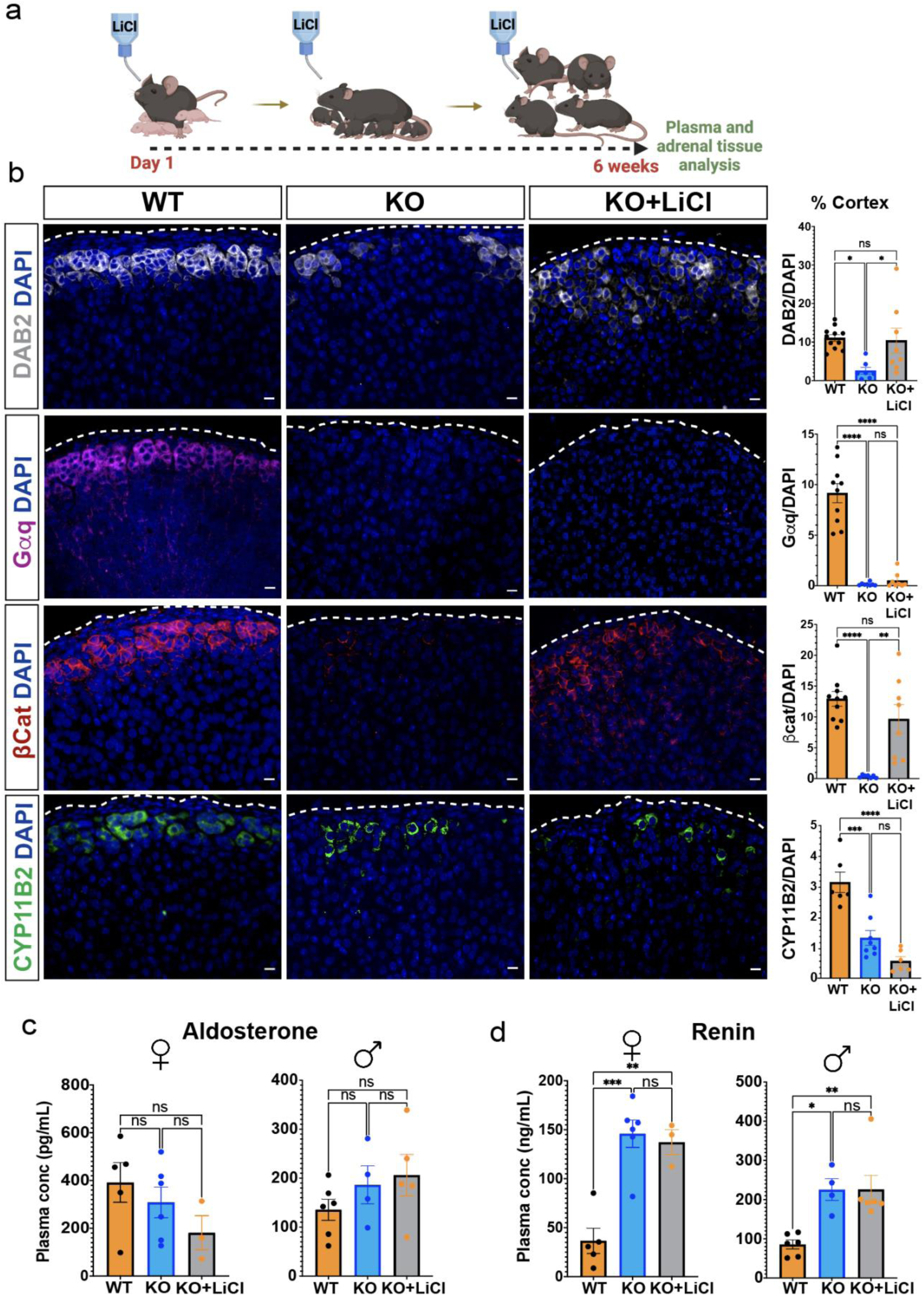
Activation of β-catenin partially rescues zG morphology, but not function, in *Wnt2b* KO mice. a. Treatment protocol with 0.06% lithium chloride (LiCl) or water from birth to 6 weeks of age. b. Representative images and quantification from 6-week-old adrenals stained for DAB2 (gray, n=11 WT, n=7 KO, n=8 KO+LiCl), Gαq (magenta, n=10 WT, n=7 KO, n=8 KO+LiCl), β-catenin (β-cat, red, n=10 WT, n=6 KO, n=8 KO+LiCl) and CYP11B2 (green, n=6 WT, n=8 KO, n=6 KO+LiCl) mice. Positive cells were quantified and normalized to nuclei (DAPI, blue) in the cortex. Scale bars: 10μm. One-way ANOVA with Tukey’s post-test. ns, not significant; *p < 0.05; **p < 0.01; ***p < 0.001; ****p < 0.0001. Data are represented as mean ± SEM. c, d. Quantification of (c) aldosterone levels from (female, n=5 WT, n=6 KO, n=3 KO+LiCl; male, n=6 WT, n=4 KO, n=5 KO+LiCl) and (d) plasma renin (female, n=5 WT, n=6 KO, n=3 KO+LiCl; male, n=6 WT, n=4 KO, n=6 KO+LiCl) mice. Data are represented as mean ± SEM. One-way ANOVA with Tukey’s post-test. ns, not significant; *p < 0.05; **p < 0.01; ***p < 0.001. Data are represented as mean ± SEM.

### Purified WNT2B does not activate canonical Wnt signaling

We recently discovered that lipid-modified WNTs (including both canonical WNT3A and non-canonical WNT5A) are released from cells via handoff from the Wntless (WLS) membrane protein to extracellular carrier proteins belonging to the Secreted Frizzled-Related Protein (SFRP) and Wnt Inhibitor Factor-1 (WIF1) families(65). In addition, we showed that both WNT3A and WNT5A are also released from cells by the ectodomain of glypicans (GPCs), an important class of WNT coreceptors(65). These results indicate that it is possible to obtain soluble WNT complexes with high stability and signaling potency. As a prelude to purifying active WNT2B complexes, we first determined whether the proteins that release WNT3A and WNT5A from Wnt-producing cells are also capable of releasing WNT2B. To test this, we stably expressed NanoLuc (NL)-tagged WNT2B in HEK293 cells and treated them with purified WNT carriers. We then quantified NL-WNT2B released into the media, as previously described for WNT3A and WNT5A(65). Notably, GPC4, SFRP2 and GPC6 were able to robustly release WNT2B from HEK293 cells (Fig. 4a), with GPC4 exhibiting the strongest effect. Due to their well-characterized functions(65), we selected GPC4 and SFRP2 for subsequent experiments.

**Figure 4.**
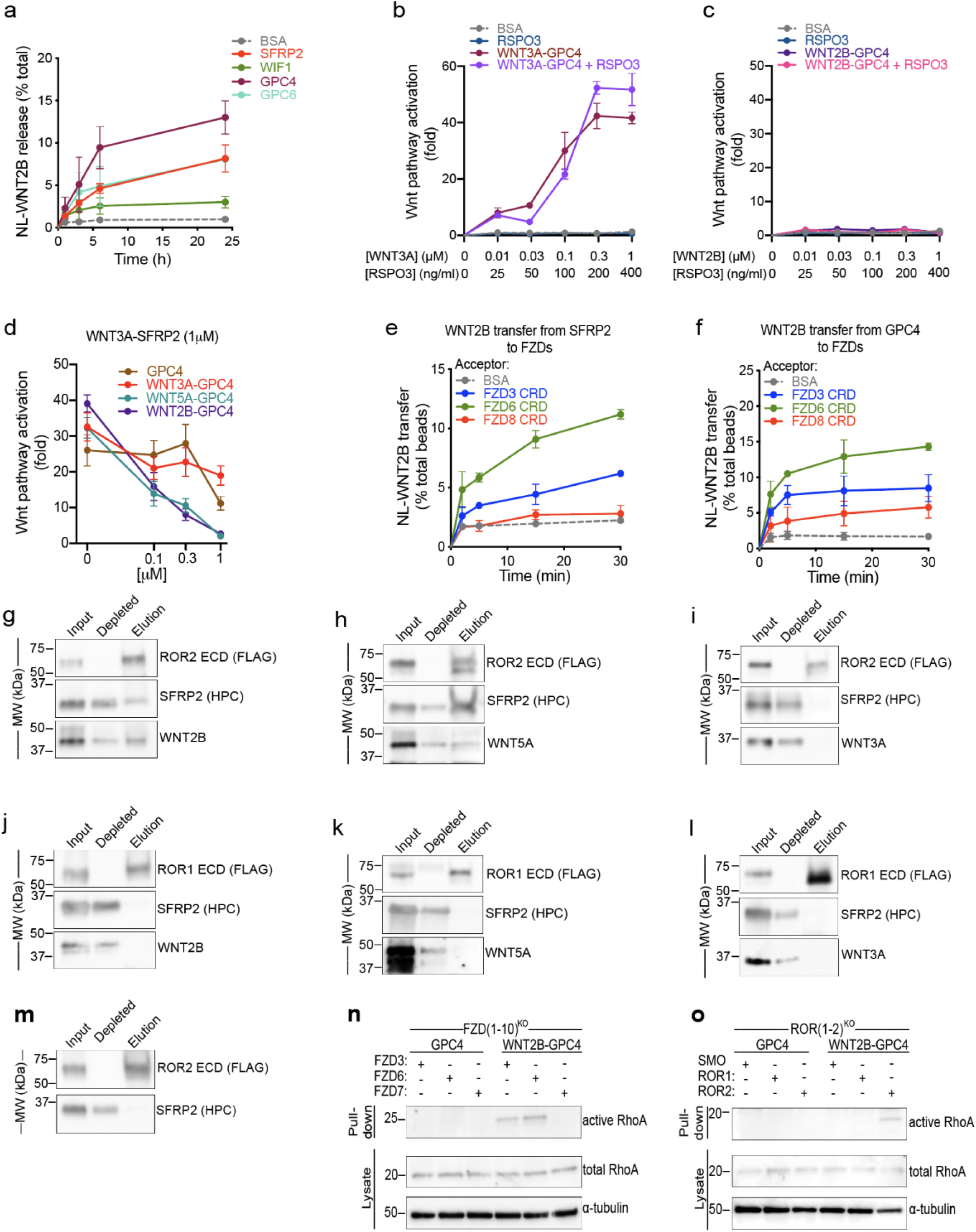
WNT2B released by SFRP2 and GPC4 activates the non-canonical Wnt/PCP pathway by binding to FZD3-CRD or FZD6-CRD, and ROR2-ECD. a. HEK293 cells stably expressing NL-WNT2B were incubated with 1μM of purified SFRP2, WIF1, GPC4 or GPC6 ectodomains in serum-free media. NL-WNT2B release was measured at various time points by NanoLuc luciferase (NL) luminescence. Bovine serum albumin (BSA) served as negative control. WNT2B is released mainly by SFRP2, GPC4 and GPC6. Data represent the mean of two biological replicates, normalized to total NL-WNT in lysates, and error bars show SD. b. R-Spondin 3 (RSPO3; 0, 25, 100, 200 and 400ng/ml) or purified WNT3A-GPC4 complex (0, 0.01, 0.03, 0.1, 0.3 and 1μM with respect to WNT3A) with or without RSPO3 (400ng/ml) was added to Wnt reporter cells. After 24h, Wnt pathway activity was measured by luciferase assay. Incubation with BSA served as negative control. RSPO3 does not potentiate WNT3A-GPC4 activity. Points represent average activation for two biological replicates, normalized to untreated cells, and error bars represent SD. See also Supplemental Figure 4a-e for protein purification and activity of WNT5A-GPC4 complex and WNT3A-carrier or WNT2B-carrier conditioned media. c. As in (b), but with purified WNT2B-GPC4 complex. WNT2B-GPC4 complex is unable to activate canonical Wnt signaling, even with RSPO3. d. As in (b), but purified WNT3A-SFRP2 complex (1μM) was mixed with varying amounts of GPC4 alone or in complex with WNT3A, WNT5A or WNT2B (0.1, 0.3 and 1μM). Both WNT5A-GPC4 and WNT2B-GPC4 complexes abolish WNT3A-SFRP2 activity, in contrast to GPC4 alone or in complex with WNT3A. See Supplemental Figure 4f for a similar experiment using WNT3A-GPC4 complex. e. NL-WNT2B-SFRP2 complex was covalently captured on HaloLink beads from conditioned media, via HT7 fused to the C-terminus of SFRP2. The beads were then incubated with purified FZD-CRDs (5µM) and NL-WNT2B release was measured at different time points by NL luminescence. Incubation with BSA (5µM) served as negative control. WNT2B is preferentially transferred to FZD3-CRD and FZD6-CRD more than FZD8-CRD. Points represent average for two biological replicates, normalized by total NL-WNT on beads, and error bars represent SD. f. As in (e), but with NL-WNT2B-GPC4 on beads. g. Purified WNT2B-SFRP2 (5μM) was incubated with the extracellular domain (ECD) of ROR2 (2.5μM), followed by immunoprecipitation with antibodies against the FLAG tag attached to ROR. Samples were analyzed by SDS-PAGE and immunoblotting. WNT2B-SFRP2 complex interacts with ROR2-ECD. See also Supplemental Figure 4g-j for protein purification and a similar experiment using purified SFRP2. h. As in (g), but with purified WNT5A-SFRP2 complex. WNT5A-SFRP2 complex binds to ROR2-ECD. i. As in (g), but with purified WNT3A-SFRP2 complex. WNT3A-SFRP2 complex does not bind to ROR2-ECD. j. As in (g), but WNT2B-SFRP2 complex (5μM) was incubated with ROR1-ECD (2.5μM). WNT2B-SFRP2 does not bind to ROR1-ECD. k. As in (j), but with WNT5A-SFRP2 complex. WNT5A-SFRP2 does not bind to ROR1-ECD. l. As in (j), but with WNT3A-SFRP2 complex. WNT3A-SFRP2 does not bind to ROR1-ECD. m. As in (g) but using SFRP2 alone. SFRP2 is unable to interact with ROR2-ECD. n. Activity of RhoA in FZD(1–10)^KO^ cells expressing FZD3, FZD6 or FZD7 was assessed by Rhotekin-RBD pull-down assay after 6h of treatment with GPC4 alone or in complex with WNT2B (2μM). RhoA endogenous levels are shown in the lysates. RhoA activity by WNT2B-GPC4, in contrast to GPC4 alone, is rescued in cells expressing FZD3 or FZD6, but not the canonical FZD7. Blotting for α-tubulin served as loading control. o. As in (n), but measuring activity of RhoA in ROR(1–2)^KO^ cells expressing ROR1 or ROR2. WNT2B-GPC4, in contrast to GPC4 alone, activates RhoA only when ROR2 expression is rescued, not ROR1. Smoothened (SMO) transfection served as negative control.

Based on the high potency of WNT3A-GPC4 complexes to activate canonical Wnt/β-catenin signaling(65), we co-expressed WNT2B with GPC4, and purified WNT2B-GPC4 complexes from conditioned media (CM) (Supplemental Figure 4a). We then tested whether WNT2B-GPC4 could trigger canonical Wnt/β-catenin signaling, using the TopFlash reporter assay(66, 67). In contrast to WNT3A-GPC4 (Fig. 4b), purified WNT2B-GPC4 did not activate canonical Wnt/β-catenin signaling, even in the presence of RSPO3 (Fig. 4c). The same result was obtained using the well-established non-canonical WNT5A(29), delivered as purified WNT5A-GPC4 (Supplemental Figure. 4b and c). Similarly, CM containing either WNT2B-carrier complexes WNT2B-SFRP2 or WNT2B-GPC4 failed to activate canonical Wnt/β-catenin signaling, in contrast to CM containing the WNT3A-carrier complexes WNT3A-SFRP2 and WNT3A-GPC4 (Supplemental Figure 4d and e).

To test whether WNT2B functions as a non-canonical WNT, we examined its ability to antagonize canonical Wnt signaling, a feature of non-canonical WNTs(68–70). Both WNT2B-GPC4 and WNT5A-GPC4 abolished canonical signaling triggered by WNT3A in a dose-dependent manner (Fig. 4d and Supplemental Figure 4f). Importantly, while excess carriers can compete with FZDs for binding to WNTs, leading to inhibition of Wnt signaling(65), increasing concentrations of WNT3A-GPC4 or GPC4 alone diminished but did not completely abolish WNT3A signaling. These results suggest that WNT2B functions as a non-canonical WNT ligand.

### WNT2B interacts with non-canonical receptors to activate the Wnt/PCP pathway via RhoA

Canonical and non-canonical WNT ligands signal through distinct FZDs based on specific coreceptor recruitment(24, 71). We have previously observed that transfer of canonical and non-canonical WNT ligands from carriers to FZDs (WNT acceptors) is specific: WNT3A is preferentially transferred to the purified extracellular cysteine-rich domain (CRD) of FZD8 (FZD8-CRD), while WNT5A is preferentially transferred to FZD3-CRD and FZD6-CRD(65). To determine if such a preference exists for WNT2B, we affinity-captured WNT2B-GPC4 and WNT2B-SFRP2 complexes to beads via a tag attached to the carrier(65), after which the beads were incubated with various purified WNT acceptors. As shown in Fig. 4e and 4f, WNT2B was rapidly transferred from SFRP2 and GPC4 to FZD3 -CRD and FZD6-CRD, but much less efficiently to FZD8-CRD. Given that FZD3 and FZD6 are established as mediators of non-canonical Wnt signaling, particularly in the Wnt/PCP pathway(72, 73), these findings are consistent with WNT2B being a non-canonical WNT ligand.

We then investigated whether WNT2B could interact with the extracellular domain (ECD) of ROR1 and ROR2 (Supplemental Figure 4g), coreceptors recognized for binding to non-canonical WNTs and promoting the activation of the Wnt/PCP signaling pathway(30, 74). Purified WNT2B-SFRP2 co-immunoprecipitated with ROR2-ECD, but not ROR1-ECD (Fig. 4g, j and Supplemental Figure 4h), similarly to the non-canonical WNT5A-SFRP2 complex (Fig. 4h, k and Supplemental Figure 4i). As expected, canonical WNT3A-SFRP2 complexes did not bind ROR2-ECD or ROR1-ECD (Fig. 4i, l). Importantly, neither ROR2-ECD nor ROR1-ECD bound the empty SFRP2 carrier, indicating a direct interaction between WNT2B and the non-canonical ROR2 coreceptor (Fig. 4m and Supplemental Figure 4j).

In vertebrates, activation of the small GTPase RhoA has been shown to be an important mediator of the non-canonical Wnt/PCP pathway(75–78). We observed that WNT2B-GPC4, like WNT5A-GPC4, activated RhoA (Supplemental Figure 4k and l). In contrast, neither purified WNT3A-GPC4 nor GPC4 alone activated RhoA. We further confirmed that WNT2B activates Wnt/PCP signaling via non-canonical receptors(72, 73), as WNT2B-GPC4 was unable to activate RhoA in human cells lacking all 10 FZD paralogs (FZD(1–10)^KO^)(65, 79), but could be rescued by expression of FZD3 or FZD6, but not FZD7, a known canonical receptor(65, 80) (Fig. 4n). Additionally, WNT2B-GPC4 failed to activate RhoA in cells lacking the ROR1 and ROR2 receptors, rescued only by expression of ROR2 (Fig. 4o). These results demonstrate that WNT2B activates Wnt/PCP signaling by activating the RhoA GTPase through non-canonical FZD receptors and the ROR2 coreceptor.

### *Wnt2b* loss disrupts the Wnt/PCP pathway in the adrenal

To further assess the role of non-canonical Wnt signaling in the adrenal cortex, we first investigated the activation of small GTPases in response to WNT2B in the mouse adrenal. Using bead-based activation assays, we demonstrated activation of RhoA(78), but not Rac1(81), another important GTPase, in WT adrenals (Fig. 5a and Supplemental Figure 5a). This suggests a specific role for RhoA in mediating Wnt/PCP signaling in the adrenal. Interestingly, a complete absence of RhoA activation was observed in *Wnt2b^-/-^* adrenals (Fig. 5a), indicating that WNT2B is required for activation of the Wnt/PCP pathway via RhoA. To extend these findings, we used bulk RNA-seq analysis to compare gene expression between *Wnt2b^-/-^* and WT adrenals. We found 1,456 differentially expressed genes between the two groups (637 up- and 819 down-regulated in *Wnt2b^-/-^*; FDR-corrected p-value < 0.05). (Supplemental Figure 5b). Gene Ontology analysis showed that several biological processes were downregulated in *Wnt2b^-/-^* adrenals, including epithelial morphogenesis, cell surface receptor signaling pathways, and positive regulation of cell adhesion (Fig. 5b). Given the role of the Wnt/PCP pathway in regulating cell polarity, adhesion, and cell rearrangement(82, 83), these findings are consistent with a critical role for WNT2B activation of the Wnt/PCP pathway in the adrenal. Furthermore, expression of genes comprising the PCP core pathway, including *Fzd3*, *Fzd6*, and *Prickle1*(73, 82, 83), as well as those involved in its regulation, such as *Cthrc1*(77) and *Dact1*(84), was reduced in *Wnt2b^-/-^* adrenals (Fig. 5c). Notably, expression of PRICKLE1, a cytoplasmic component of the non-canonical Wnt signaling pathway that establishes planar cell polarity(82), was predominantly observed in the zG of both mouse and human adrenals but was absent in *Wnt2b^-/-^* adrenals (Fig. 5d and Supplemental Figure 5c). Moreover, expression of genes encoding core components of the PCP pathway remained unaffected in a mouse model of zG-specific β-catenin LOF(3), in which we observed a marked decrease in canonical Wnt/β-catenin signaling, indicating the independence of PCP core protein regulation from canonical Wnt/β-catenin signaling (Supplemental Figure 5d). Together, these results imply that Wnt/PCP signaling is active in the zG and is strongly disrupted by loss of *Wnt2b*. This supports the hypothesis that WNT2B, produced by the adrenal capsule, regulates morphogenesis of the underlying zG through activation of the Wnt/PCP signaling pathway and maintenance of PCP components.

**Figure 5.**
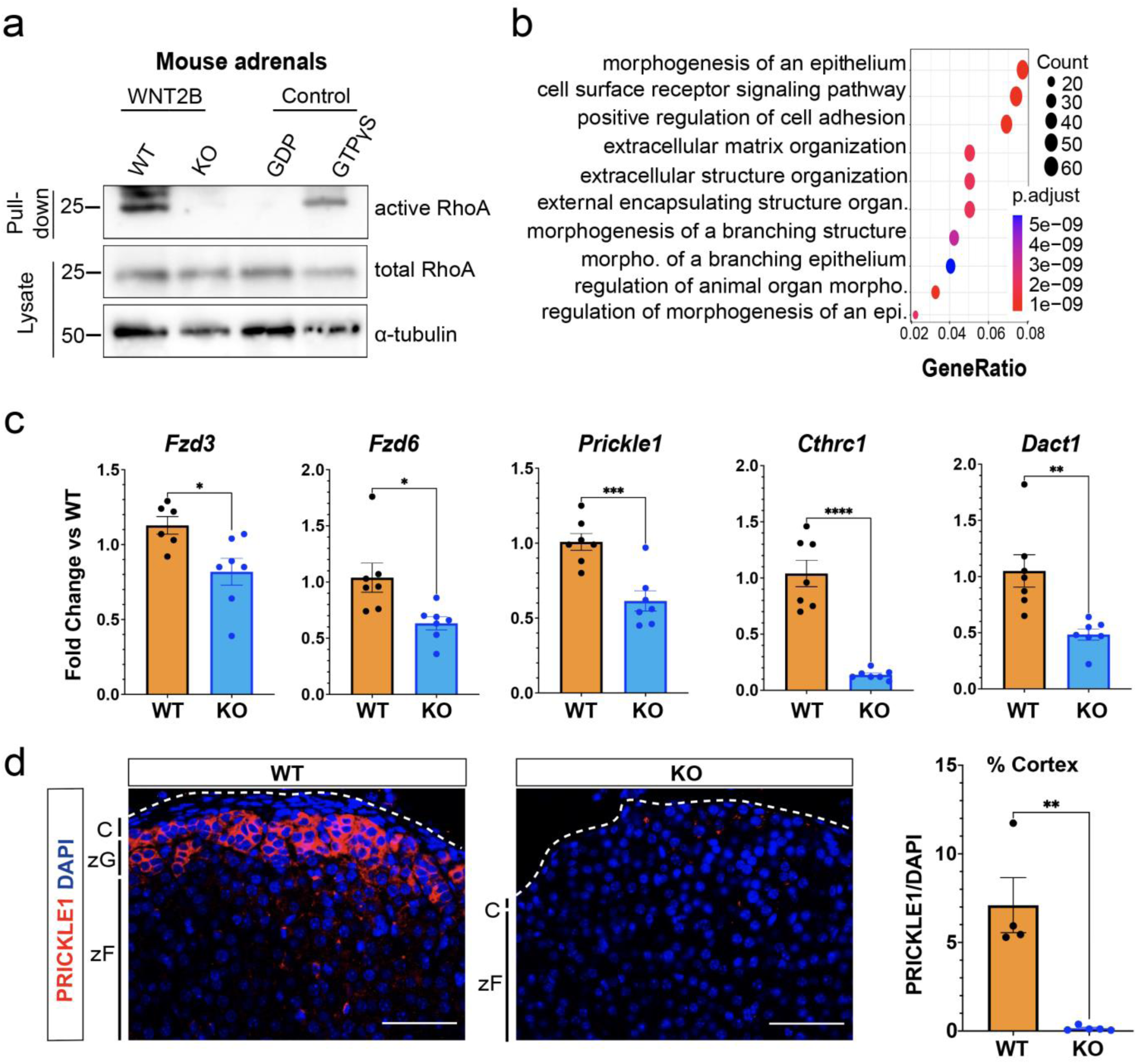
WNT2B deficiency disrupts Wnt/PCP signaling in the adrenal. a. Activity of RhoA in WT and KO adrenals was assessed by Rhotekin-RBD pull-down assay using adrenal lysates. GTPγS and GDP treated adrenal lysates served as positive and negative controls, respectively. Total RhoA and α-tubulin served as loading controls. b. Dot plot depicting Gene Ontology (GO) Gene Set enrichment analysis of genes downregulated in KO vs WT. c. QRT-PCR was performed in WT and KO adrenals for *Fzd3* (n=6 WT, n=7 KO), *Fzd6* (n=7 WT, n=7 KO), *Prickle1* (n=7 WT, n=7 KO), *Cthrc1* (n=7 WT, n=7 KO) and *Dact1* (n=7 WT, n=7 KO) from female mice. Two-tailed Student’s t-test. *p<0.05; **p < 0.01 ***p < 0.001; ***p < 0.0001. Data are represented as mean ± SEM. d. Representative images and quantification from adrenals stained for PRICKLE1 (red, n=4 WT, n=5 KO). Positive cells were quantified and normalized to nuclei (DAPI, blue) in the cortex. Scale bars: 50μm. Two - tailed Student’s t-test. **p < 0.01. Data are represented as mean ± SEM. C, capsule; zG, zona glomerulosa; zF, zona fasciculata.

### Components of Wnt/PCP signaling are conserved in mouse and human adrenal

To assess whether mediators of non-canonical Wnt signaling in the adrenal cortex are conserved in humans and mice, we analyzed single nuclei RNA sequencing (snRNA-seq). Consistent with our initial findings (Fig. 1a), *WNT2B* was localized to the SHH-responsive GLI1+ RSPO3+(41, 55) capsular cells in both human and mouse adrenals (Fig. 6a-b, Supplemental Figure 6a-b). In addition, the non-canonical WNT receptors *FZD3*, *FZD6*, and the *ROR2* coreceptor were expressed in the zG in both human and mouse adrenals (Fig. 6c-d). These findings were further validated using RNAscope on human and mouse adrenals (Fig. 6e-f). This collective evidence underscores the conservation across species of Wnt/PCP signaling components in the adrenal cortex and points to a potential role for WNT2B in the human adrenal.

**Figure 6.**
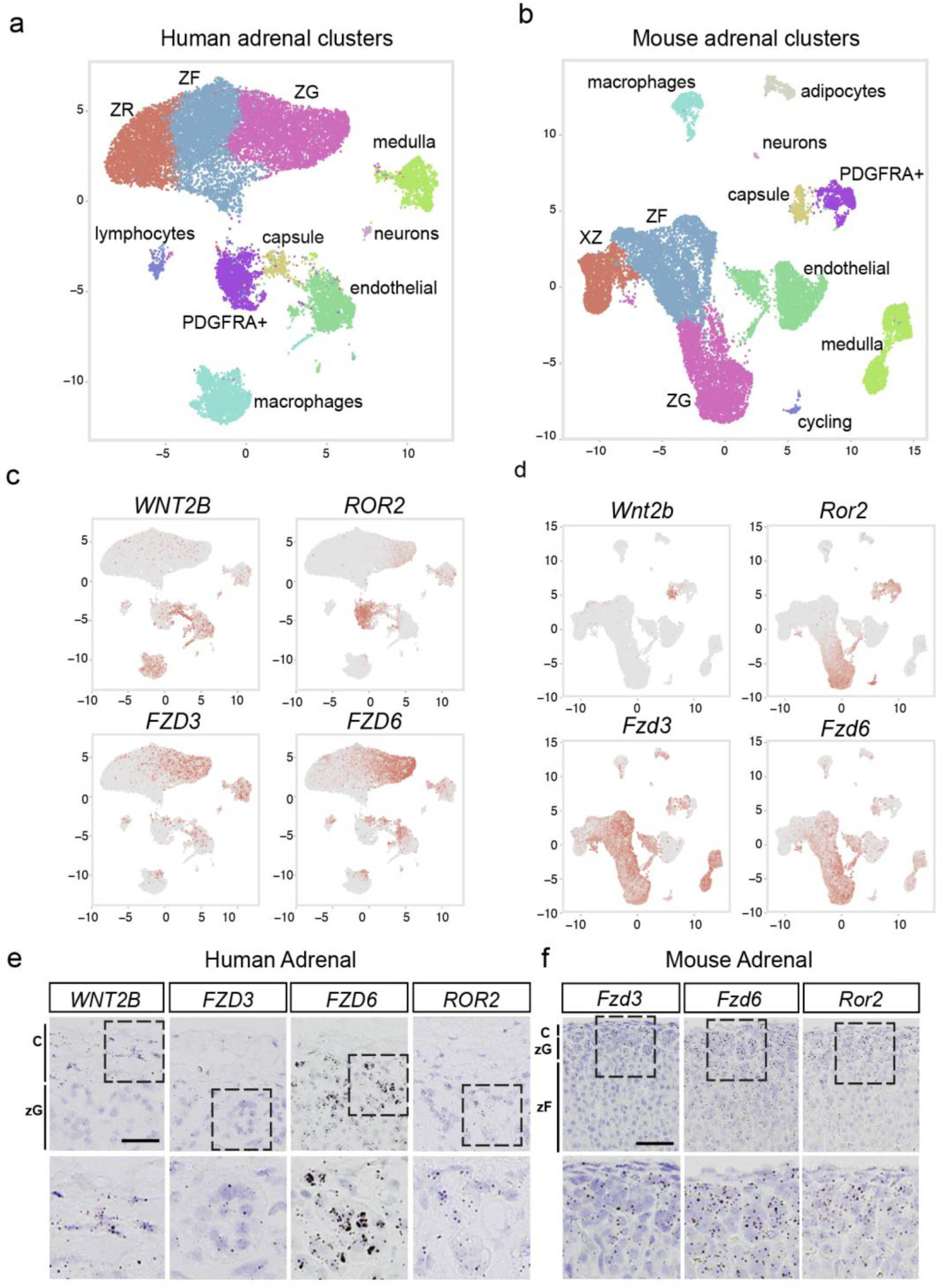
Components of Wnt/PCP signaling are conserved across mouse and human adrenals. UMAP plots of snRNAseq from human (a) and mouse (b) adrenals depicting similarly diverse cell types including cortical and non-cortical cells. Expression patterns of *WNT2B*, *ROR2*, *FZD3* and *FZD6* projected over the UMAP projections from human (c), and mouse (d) adrenals. e. Representative smISH images of *WNT2B*, *FZD3*, *FZD6* and *ROR2* expression in the adrenal cortex of human adrenals. Scale bar: 25μm f. Representative smISH images of *Fzd3*, *Fzd6* and *Ror2* expression in the adrenal cortex of mouse adrenals. Scale bar: 25μm

### Homozygous loss of *WNT2B* results in congenital hypoaldosteronism in humans

Finally, to investigate the role of WNT2B in aldosterone production in humans, we analyzed a rare cohort of three individuals with WNT2B deficiency (Table 1). Individuals A and B were siblings and carried homozygous LOF mutations in *WNT2B*(85), while individual C carried compound heterozygous LOF mutations(86). All three individuals exhibited Congenital Diarrhea and Enteropathy (CoDE) syndrome requiring parenteral nutrition to achieve euvolemia. Analysis of RAAS activity revealed elevations in plasma renin concentrations (or plasma renin activity) with compensated plasma aldosterone levels, resulting in low aldosterone/renin ratios (ARR). Individual C also received a trial of fludrocortisone, a synthetic steroid with high mineralocorticoid activity, revealing mineralocorticoid sensitivity consistent with intact mineralocorticoid receptor function (Table 1). These results indicate that WNT2B is essential for aldosterone production in humans and that WNT2B deficiency leads to a newly identified form of Familial Hyperreninemic Hypoaldosteronism, designated here as Type 4.

**Table 1.**
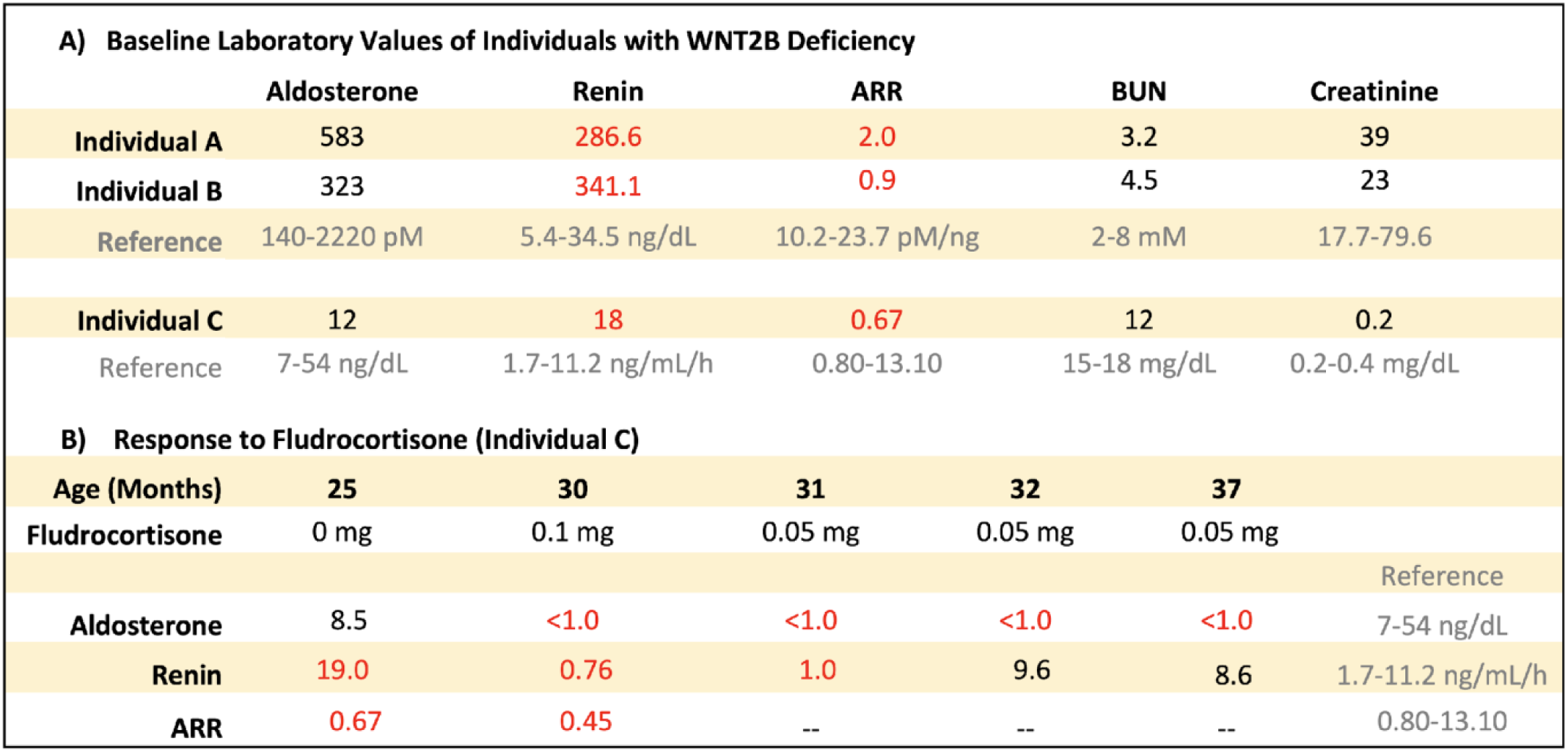

## Discussion

In this study, we demonstrate that *WNT2B* is expressed in the adrenal capsule of both mice and humans, and that loss of WNT2B results in adrenal hypoplasia, disruption of zG morphology, and a reduction in aldosterone-producing cells. Remarkably, our data reveal that WNT2B deficiency results in Hypo-A in both mice and humans. Additionally, we replicate prior findings that a common non-coding variant in *WNT2B* is associated with increased risk of PA(46, 47). Although connecting non-coding variant signals from GWAS to the genes mediating their effects is often challenging, our data provide compelling evidence from both mice and humans that this variant acts through WNT2B. Furthermore, we elucidated that WNT2B functions as a non-canonical WNT ligand, activating the Wnt/PCP signaling pathway within the adrenal cortex. Collectively, our results establish WNT2B as a crucial non-canonical WNT ligand essential for zG formation, maintenance, and aldosterone production.

The adrenal capsule, a thin layer of mesenchymal cells situated as the outermost compartment of the adrenal gland, plays a vital role in communicating with the subcapsular zG to facilitate homeostatic cellular renewal and to maintain the distinctive properties of the zG(33, 41, 55, 56). We show that *WNT2B* is expressed in the capsule and that the loss of *Wnt2b* leads to a reduction in adrenal size, likely due to a near-complete loss of the histological zG. This was supported by decreased expression of established zG markers such as β-catenin, DAB2, Gαq, and CYP11B2 in *Wnt2b^-/-^* adrenals, which are critical for zG differentiation and function. Moreover, conditional KO of *Wnt2b* in GLI1+ RSPO3+ capsular cells of the adult mouse leads to similar defects in zG morphology, as reflected by downregulation of CYP11B2 and DAB2 expression. Further analysis of adrenal cells from *Wnt2b^-/-^* mice revealed a marked reduction in the expression of *Shh,* a key marker associated with adrenal progenitor cells in the zG, and its target *Gli1*, which is expressed in the capsule(55, 56). Downregulation of these genes was accompanied by a thinning of the capsule in the *Wnt2b^-/-^* adrenals. These findings are consistent with mouse models where genetic loss of *Shh* results in hypoplastic adrenals with a thinner capsule(56, 57) and underscore the importance of cortex-to-capsular SHH signaling as a mechanism for homeostatic adrenocortical renewal. Taken together, these data show that WNT2B, produced in the adrenal capsule, is essential for the proper development and maintenance of the underlying zG in the adrenal gland.

The primary function of the zG is production and secretion of aldosterone into the bloodstream as an integral part of the RAAS(5). Despite a significant scarcity of aldosterone-producing cells in the adrenal cortex of *Wnt2b^-/-^* mice, plasma aldosterone levels were surprisingly observed to be within the normal range. This coincided with a marked increase in plasma renin levels, indicating activation of the RAAS to compensate for the lower number of aldosterone-producing cells. Further analysis of adrenal slices *ex vivo*, without compensatory physiological mechanisms, revealed impaired aldosterone production due to WNT2B deficiency. Together, our data indicate that *Wnt2b^-/-^* mice, in response to physiological demands (e.g., volume depletion), increase renin levels to maintain inappropriately normal aldosterone levels, despite the reduced number of aldosterone-producing cells.

Canonical Wnt/β-catenin signaling has been shown to regulate zG differentiation and aldosterone production in the adrenal cortex(8, 31, 34, 87). Surprisingly, when *Wnt2b^-/-^* mice were treated with LiCl, an activator of the canonical Wnt/β-catenin pathway(62, 63), we did not observe a decrease in plasma renin levels, likely due to the failure to induce Gαq and CYP11B2 in the zG. Despite it having been described as a canonical WNT ligand(50, 88–92), WNT2B did not activate the canonical Wnt/β-catenin signaling pathway. In stark contrast, WNT2B exhibited a distinctive feature of non-canonical WNTs: the ability to antagonize canonical Wnt signaling(68–70). Further analysis revealed that WNT2B activates the Wnt/PCP pathway through the non-canonical WNT receptors FZD3 and FZD6 and the ROR2 coreceptor(72–74). This was supported by WNT2B’s ability to activate the GTPase RhoA, a mediator of the Wnt/PCP pathway(76, 78). Furthermore, our findings revealed that RhoA is activated in WT adrenals, likely contributing to both zG morphogenesis and aldosterone production. In contrast, activated RhoA was not detected in W*nt2b*^-/-^ adrenals. These results establish that WNT2B functions as a non-canonical WNT ligand, primarily activating the Wnt/PCP signaling pathway in the adrenal.

Wnt/PCP signaling is a highly conserved pathway essential for coordinating cell polarity and morphogenesis across multiple tissues and is especially important for rosette formation(78, 82, 83, 93). Notably, rosettes are a multicellular structure essential for postnatal zG development and are a hallmark of aldosterone-producing cell clusters in mice and humans(2, 3), which are essentially absent from *Wnt2b^-/-^* adrenals. The Wnt/PCP pathway relies on the asymmetric distribution of core PCP components, which in mammals includes the orthologues of *Drosophila melanogaster* proteins: FZD3/6, Dishevelled (DVL1-3), Van Gogh (VANGL1/2), Flamingo (CELSR1-3), Prickle (PK1-2), and Diego (ANKRD6)(94). Generally, these core PCP signaling molecules interact both across cell membranes and intracellularly to establish two complexes on opposing sides of each cell(82, 83). The conserved expression of *FZD3*, *FZD6*, and *ROR2* within the zG of both humans and mice indicates that this structure possesses the necessary components for activation of the Wnt/PCP pathway. This also implies that WNT2B, produced and secreted by GLI1+ RSPO3+ capsular cells (WNT2B-producing cells), are transferred to the zG cells (WNT2B-receiving cells), possibly establishing a gradient, to initiate Wnt/PCP signaling. In support of this model, we found that another core PCP protein, PRICKLE1(82), is expressed in the zG and is reduced in *Wnt2b^-/-^* mice. Importantly, these factors appear to function independently of the canonical Wnt/β-Catenin pathway in the zG. How WNT2B interacts with other regulatory mechanisms to mediate Wnt/PCP signaling during zG formation, maintenance, and function remains to be fully elucidated.

FHH refers to a group of inherited disorders characterized by abnormally high levels of renin in the blood (hyperreninemia) and low levels of aldosterone hormone (hypoaldosteronism)(6). FHH Type 1 is caused by mutations in the *CYP11B2* gene(9, 95), while type 2 is associated with unknown mutations not linked to *CYP11B2*(7). Recently, LOF mutations in the R-spondin receptor *LGR4/GPR48* have been implicated in abnormal zG differentiation and FHH(8). In our study, we assessed the RAAS in three individuals with confirmed *WNT2B* deficiency(85, 86). We observed low-to-normal aldosterone levels and elevated renin levels, resulting in a low aldosterone/renin ratio. These findings indicate that WNT2B deficiency represents a new form of FHH, designated here as Type 4. Notably, these findings contrast with the non-coding variant (rs3790604) in *WNT2B*, which is associated with a predisposition to PA(46, 47). Further investigation is needed to determine if this allele leads to an increase in the number of aldosterone-producing cells or simply an increase in aldosterone production by the zG.

In conclusion, this study provides significant insights into the role of WNT2B in adrenal gland development and function. Our findings demonstrate the importance of paracrine signals in maintaining the integrity of the adrenal cortex, with WNT2B produced by the adrenal capsule playing a crucial role in zG formation, maintenance, and aldosterone production. We demonstrate that WNT2B functions as a non-canonical WNT ligand, activating the Wnt/PCP signaling pathway in the adrenal gland. Furthermore, we identified WNT2B deficiency as a new form of familial hyperaldosteronism (FHH), designated here as Type 4, which implies that the common non-coding variant in *WNT2B* associated with increased susceptibility to PA involves a GOF mechanism. These results highlight the complex interplay between paracrine signals and cell populations in regulating endocrine function and the role of both canonical and non-canonical Wnt pathways within the adrenal gland. This study provides valuable insights into the mechanisms underlying adrenal homeostasis and identifies potential therapeutic targets for the treatment of adrenal disorders.

## ACKNOWLEDGEMENTS

We gratefully acknowledge the clinical subjects and the *All of Us* participants for their contributions, without whom this research would not have been possible. We also thank the National Institutes of Health’s *All of Us* Research Program for making available the participant data examined in this study. This work was supported by a Physician-Scientist Career Development Award from K12DK133995 (to SA), R01DK123694 (to DTB), R01HL155834 (to AFT), R01DK062027 to (GDH), R01GM122920-05 and R35GM153357-01 (to AS) and Cell Biology Education and Goldberg Fellowship Fund (to TdAM). We thank members of the Salic, Breault and Hammer laboratories for constructive comments and ongoing support.

## AUTHOR CONTRIBUTION

KSB, DWL III, TdAM, AS, DTB, and GDH conceptualized the project and designed the analysis. KSB designed, performed and analyzed the majority of the experiments. DWL III designed and conducted key experiments, particularly those involving the inducible CRE mouse model, snRNA-seq and RNAscope. TdAM performed all mechanistic studies related to Wnt signaling *in vitro*. KSB, DWL III, TdAM, DLC, AS, DTB, and GDH contributed to drafting the manuscript. KSB, DWL III, TdAM, CR, TD, CL, KJB, NAG, PQB, MB, AEO, AFT, AML, DRM, WR, and DLC performed experiments and analyses. NAG and PQB conducted the *ex vivo* adrenal experiment and analysis. DTB and SA carried out patient analysis and disease definition. DRM, DWL III and AML performed snRNA-seq experiments and analyses. AS and JNH were responsible for the GWAS analysis. All authors participated in data interpretation and critically reviewed the manuscript. The first co-authorship order was determined through a collaborative discussion, considering the significance and scope of each author’s contributions to the research.

## DATA ACCESS STATEMENT

This study used data from the *All of Us* Research Program’s Controlled Tier Dataset v7.1, available to authorized users on the Researcher Workbench.

## Methods

### Sex as a biological variable

Our study examined both male and female human subjects, as well as male and female mice, and found similar results across both sexes.

### Mice

Experiments involving *Wnt2b* global KO (*Wnt2b^-/-^*) mice were carried out in accordance with protocols approved by the Boston Children’s Hospital’s Institutional Animal Care. *Wnt2b^fl/fl^* mice (a generous gift from T. Yamaguchi, NCI/NIH(52)) were crossed with *CMV-Cre* mice (Jackson labs) to generate *Wnt2b^fl/-^* mice, which were then intercrossed to generate *Wnt2b^-/-^* mice. Male and female mice were used for experiments at ∼two months of age and *Wnt2b^+/+^* (wild type, WT) littermates were used as controls.

Experiments involving LiCl rescue were carried out with *Wnt2b^-/-^* mice treated with 0.06% lithium chloride (LiCl) in drinking water, as previously reported(62, 63). Mice used for experiments received LiCl through the mother’s breast milk for the first three weeks of life and from their own LiCl-treated water source for the following three weeks (until 6 weeks of age).

The *AS^Cre/+^ :: Ctnnb1^fl/fl^* mouse strain has been described previously(3). Mice were studied at 3 months of age and *AS^Cre/+^*mice were used as controls.

Experiments involving conditional *Wnt2b* cKO (*Gli1^CreER/+^* :: *Wnt2b^fl/fl^*) mice were carried out in accordance with protocols approved by the University Committee on Use and Care of Animals at the University of Michigan. *Wnt2b^fl/fl^* mice (a generous gift from T. Yamaguchi, NCI/NIH(52)) were crossed with CAG-flpo mice(96) (Jackson labs) to remove the NeoR cassette from the original floxed *Wnt2b* allele. To generate *Wnt2b* cKO mice targeting the adrenal capsule, *Wnt2b^fl/fl^* mice (minus the NeoR cassette) were crossed with *Gli1^CreERT2/+^* mice(59) to generate *Wnt2b* cKO mice. Male and female mice were used for experiments at six-seven weeks of age and *Wnt2b^fl/+^* and *Wnt2b^+/+^* mice were used as controls. *Wnt2b* cKO mice were injected with tamoxifen (Sigma-Aldrich), dissolved in 10% ethanol and 90% corn oil to a final concentration of 10mg/ml daily, for five consecutive days (IP 1mg/20g body weight). Adrenals were harvested for IHC and measurement of RNA expression four weeks following tamoxifen injection.

All mice were maintained on a mixed background under a 12-hour light/dark cycle with *ad lib.* access to food and water.

### Adrenal dissection and preparation

After dissection, adrenals were cleaned of periadrenal fat, rinsed in phosphate buffered saline (PBS), and weighed. For immunohistochemistry, adrenals were fixed in 4% paraformaldehyde (PFA) at 4°C overnight. Adrenal weights were normalized to mouse body weights, which were obtained one day prior to sacrifice to minimize induction of the stress response. Processed adrenals were paraffin-embedded and cut in 5 µm sections for histological use.

### Immunostaining

Sections were rinsed in xylene, an ethanol gradient and then PBS. Antigen retrieval was performed in Tris-EDTA pH 9.0. Sections were blocked in 5% Normal Goat Serum in PBS for 1h at RT. Primary antibodies were diluted 1:200 in 5% NGS in PBS and incubated on sections at 4°C overnight. Slides were washed three times for 5 min in 0.1% Tween-20 in PBS. Secondary antibodies were diluted in 1:300 in PBS and incubated on sections at RT for 1–2 h. For nuclear staining, DAPI (4′,6-diamidino-2-phenylindole) was added to secondary antibody mixture at a final concentration of 1:1000. After three 5-min washes with 0.1% Tween-20 in PBS, slides were mounted with ProLong Gold Antifade Mountant (Thermo Fisher Scientific, P36930). Primary antibodies used for this application include: Mouse anti-β-catenin (BD Biosciences, 610153), Rabbit anti-CYP11B2 and anti-CYP11B1 (kindly provided by Dr. Celso E. Gomez-Sanchez), Mouse anti-Dab2 (BD Biosciences, 610464), Rabbit ant-Dab2 (Cell signaling, 12906), Rabbit anti-Gαq (Abcam, ab75825), Rabbit anti-Prickle1 (Proteintech, 22589-1), Rabbit anti-Lef1 (Abcam, ab137872), Mouse anti-NF2R2 (R&D, PP-H7147-00) and Rabbit anti-Akr1b7 (kindly provided by Dr. Pierre Val and Dr. Antoine Martinez). The following secondary antibodies were used: Alexa Fluor 647-conjugated goat anti-rabbit IgG, Alexa Fluor 594-conjugated goat anti-mouse IgG (Invitrogen).

### Immunofluorescence quantification

Images were acquired using a Nikon upright Eclipse 90i microscope. For each image, three Z-stacks were collected and deconvoluted to achieve the best resolution using the LIS-Elements Nikon software. Single adrenal images were stitched together and adjusted for brightness and contrast using ImageJ software. Brightness levels were optimized to enhance visibility without causing overexposure of pixel data, and regions with paraffin folding or nonspecific background were removed using ImageJ. Additionally, medulla and nonspecific staining above the capsule were removed, keeping only the entire adrenal cortex. Whole adrenal images were exported in PNG file format (separate file for each channel) and imported into Photoshop (version 25.2.0) as separate layers. Quantification of positive areas in the adrenal cortex for β-catenin, DAB2, Gαq, CYP11B2, PRICKLE1 and NF2R2 was conducted using one complete equatorial section per mouse adrenal gland. This was achieved using Photoshop’s color selection tool to select the stained positive regions. Quantification was based on pixel count/area for the positive regions using the histogram. Specifically, the pixel area within stain-positive regions was measured and normalized to the pixel count/area of DAPI staining to control for variations in cell number. Normalization of the positive areas to DAPI staining ensures accurate comparison and interpretation of results across different samples.

### Floating section immunofluorescence

After fixation, adrenals were sectioned using a vibratome as described previously(3). The 100 µm floating adrenal sections were incubated with 1:100 diluted rat anti-Laminin β1 (Santa Cruz, sc-33709) and 1:200 diluted rabbit DAB2 (Cell Signaling Technologies, 12906) primary antibodies overnight at 4°C. Secondary antibodies, Alexa Fluor 488-conjugated goat anti-rabbit IgG and Alexa Fluor 647 conjugated goat anti-rat IgG (Invitrogen), were used at a diluted of 1/200. Imaging was performed using a Zeiss LSM 510 confocal microscope (Carl Zeiss AG) equipped with a 40X/1.3 oil immersion PLAN-APOCHROMAT objective. DAB2 labeling was used to image the zG of adrenal slices. For quantification, the number of DAB2+ cells in glomerular structures delineated by Laminin β1 labeling were counted manually. A zG rosette was defined as a Laminin β1-encircled glomerular structure containing five or more DAB2+ cells. At least three different regions of each adrenal were imaged and quantified. Each dot in the graph represents the average zG-rosette number per area for one animal.

### Single molecule *in situ* hybridization and quantification

For single molecule *in situ* hybridization experiments, adrenals were fixed in 10% neutral buffered formalin (NBF FischerBrand, #427-098) for 24 h at room temperature. All smISH tissue preparation and experiments were done in RNase-free conditions. Adrenal sections used in smISH experiments were stored in sealed slide boxes with desiccant (Sorbent Systems, U1MSNWP) and used within one week of sectioning. Single molecule ISH was performed using RNAScope 2.5 HD Brown Detection Kit (Advanced Cell Diagnostics, abb. ACD Cat# 322310) according to manufacturer’s instructions. Target retrieval was performed for 7 minutes, as previously reported (Cat# 322000). Slides were counterstained with 50% Gill’s hematoxylin (Millipore Sigma, GSH132-1L) and mounted with EcoMount (Biocare Medical, EM897L). The following probes were used: Mm-Wnt2b (ACD, 405031), Mm-Shh (ACD, 314361), Mm-Cyp11b2 (ACD, 505851), Mm-Wnt4 (ACD, 401101). Negative control (Mm-Dapb, ACD, 310043) and positive control (Mm-Polr2a, ACD, 312471) probes were used for each experiment to verify sample quality. 40X images were obtained on a Nikon E800 microscope and analyzed in ImageJ. At least 3 technical replicate images per biological replicate was reported.

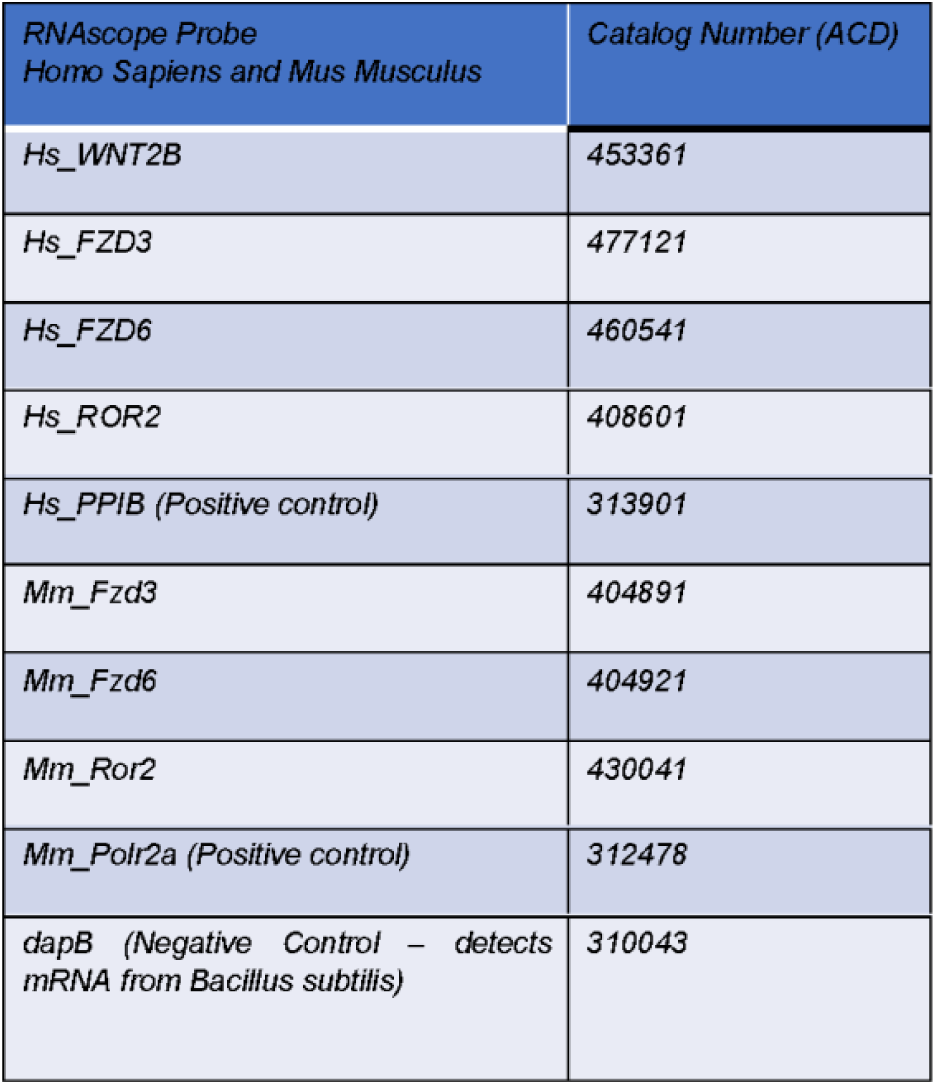

| RNAscope Probe<br>Homo Sapiens and Mus Musculus | Catalog Number (ACD) |
| --- | --- |
| Hs_WNT2B | 453361 |
| Hs_FZD3 | 477121 |
| Hs_FZD6 | 460541 |
| Hs_ROR2 | 408601 |
| Hs_PPIB (Positive control) | 313901 |
| Mm_Fzd3 | 404891 |
| Mm_Fzd6 | 404921 |
| Mm_Ror2 | 430041 |
| Mm_Polr2a (Positive control) | 312478 |
| dapB (Negative Control – detects<br>mRNA from *Bacillus subtilis*) | 310043 |

### Gene expression analysis

Total RNA was purified from *Wnt2b^-/-^* and WT whole adrenals cleaned of periadrenal fat and homogenized in TRI®Reagent (Sigma) using the Direct-zol™ RNA kit (Zymo Research), following the manufacturer’s protocol. Further processing of total RNA involved reverse transcription into cDNA using the High-Capacity cDNA Reverse Transcription Kit (Life Technologies). Gene expression analysis was performed by real-time quantitative PCR (qPCR) using the QuantStudio 6 Flex thermocycler (Life Technologies). Technical duplicates were used to control for technical variability. The TaqMan Universal PCR Master Mix and the following mouse Taqman primers from Life Technologies were used: *Wnt4* (Mm01194003_m1), *Wnt2b* (Mm00437330_m1), *Cyp11b2* (Mm01204955_g1), *Shh* (Mm00436528_m1), *Actb* (Mm02619580_g1), *Lef1* (Mm00550265_m1), *Dab2* (Mm01307290_m1) and *Gli1* (Mm00494654_m1). *Actb* transcripts, encoding β-actin, were used as the internal control and data were expressed using the 2−ddCt method97. Total RNA from cKO adrenals was analyzed by qPCR using Power SYBR Green PCR Master Mix (Invitrogen) on a QuantStudio 3 thermocycler. qPCR primers were as follows: mouse Cyp11b2 F 5’-GCACCAGGTGGAGAGTATGC-3’, R 5’-CCATTCTGGCCCATTTAGC-3’; mouse Wnt2b F 5’-CATGCTCAGAAGCAGCCGGG-3’, R 5’-GTTGATCATGGTGCCGACCG-3’; Actb F 5’-GTGACGTTGACATCCGTAAAGA-3’, R 5’-GCCGGACTCATCGTACTCC-3.

### Steroid Measurements

Plasma from mice was obtained through retro-orbital blood collection using sodium heparin-coated evacuated tubes (Fisher Scientific)(34). The collected blood was centrifuged at 1800 x g for 15 minutes at 4°C to separate plasma and plasma samples were stored at -80°C until further analysis. Aldosterone and corticosterone were quantified by liquid chromatography-tandem mass spectrometry (LC-MS/MS), as previously described(97).

### Mouse renin assay

Plasma renin from mice was obtained through retro-orbital blood collection using sodium heparin-coated evacuated tubes (Fisher Scientific)(34). The collected blood was centrifuged at 1800 x g for 15 minutes at 4°C to separate plasma and plasma samples were stored at -80°C until further analysis. Samples were thawed and analyzed using Mouse Renin ELISA Kit (Thermo Fisher, EMREN1) according to manufacturer’s instructions. Plasma samples were diluted at 1:15 for use as previously described(64).

### *Ex vivo* Aldosterone Secretion Assay

Aldosterone production from mouse adrenal slices was measured as previously described(61). Briefly, adrenal glands were harvested from 6-8-month-old male WT and Wnt2b^-/-^ mice, embedded in 3.2% agar/PIPES buffer and sectioned on a vibratome (60–70 µm slices, Microslicer Zero-1, Ted Pella) in ice-cold PIPES incubation buffer (in mM: 20 PIPES, 140 NaCl, 3 KCl, 1 CaCl2, 1 MgCl2, 25 D-Glucose, 5 NaHCO3, pH 7.3 [adjusted with 10N NaOH]). Slices were laid flat on cell culture inserts (Millicell, PICM03050) in 6-well plates (Corning, Costar 3513) and incubated in cell culture media (DMEM/Nutrient mixture F-12 Ham powder, Millipore Sigma, D9785) at 37°C with 5% CO2. After a 2-hour equilibration period, the slices were transferred to fresh media for 30 minutes and media was collected for Aldosterone (RIA, Tecan US, c., MG13051) and Corticosterone (ELISA, R&D Systems, KGE009) assays.

### Bulk RNA sequencing

Adrenals used for RNA sequencing were dissected as described above, added to Trizol, and stored at -80°C until use. RNEasy Mini Kit (Qiagen) reagents were used to isolate total RNA. Frozen adrenals were added to 600µL Buffer RLT (Qiagen) with 2-mercaptoethanol (10 uL/mL) in sterile Lysing Matrix D tubes (MP Biomedicals, 6913100) and homogenized 2 x 30 seconds using a Bead Bug Homogenizer. Extracted RNA was eluted in 20µL of RNase-free deionized water and measured for concentration and quality on a NanoDrop spectrophotometer. Library prep and next-generation sequencing was carried out as previously decribed(3).

For data analysis, sequencing metrics such as base quality score and number of sequences were assessed with FastQC (version 0.11.9) (https://www.bioinformatics.babraham.ac.uk/projects/fastqc/). Read adapters were trimmed with the bbduk tool from bbtools (version 38.96) (https://sourceforge.net/projects/bbmap/). Paired-end sequences were aligned to the mouse transcriptome reference sequence (release M38, obtained from gencodegenes.org) using kallisto (version 0.46.2)(98). Downstream analyses were performed in R using Bioconductor tools. Expression values were summarized at the gene level using the lengthScaledTPM method from tximport (version 1.18.0)(99). Inter-experiment gene-level expression values were scaled to library size using the TMM method from edgeR (version 3.32.0)(100). Unwanted and hidden sources of variation were removed from the data using sva (version 3.46.0)(101). Differential gene expression analysis was performed with limma (version 3.46.0)(102). Heatmaps and volcano plots were built with pheatmap and EnhancedVolcano, respectively (Kolde, R. pheatmap: Pretty Heatmaps. R package v.1.0.12 https://CRAN.R-project.org/package=pheatmap (2019); Blighe, K., Rana, S. & Lewis, M. EnhancedVolcano: publication-ready volcano plots with enhanced coloring and labeling. R package v.1.12.0 https://github.com/kevinblighe/ EnhancedVolcano (2021)). Gene ontology enrichment analysis of the differential expressed genes was performed with the enrichGO function from clusterProfiler (version 3.17.5)(103).

### Single Nuclei RNA sequencing analysis

Data from human (experiments ENCSR362YDM and ENCSR724KET, 26-year-old male and 16-year-old female subjects, respectively) and mouse (60-day old; experiments ENCSR356VJZ, ENCSR908CQZ, ENCSR244OUG and ENCSR749GDE) single-nuclei RNA-seq were downloaded from ENCODE and processed. Raw sequencing data were aligned to the human and mouse reference genomes (contigs GRCh38 and GRCm39, respectively) as appropriate and quantified using cellranger-arc count (version 2.0.2). Output files containing gene expression count data were processed in R using Seurat (version 5.0.1)(104). Low-quality nuclei (i.e., cells exhibiting a high percentage of mitochondrial genes and a low number of features) were flagged and removed using miQC (version 1.10.0)(105). Doublets were identified and removed using DoubletFinder (version 2.0.4)(106). Experiments from human and mouse datasets were normalized using the SCTransform function from Seurat. The percentage of reads mapping to mitochondrial genes were used as covariates to regress against during the normalization process. For visualization purposes, imputation of missing values was performed with alra (from SeuratWrappers package version 0.3.2)(107). Human and mouse datasets were integrated using harmony (version 1.2.0)(108). Clusters were identified using the FindClusters function from Seurat using the Leiden algorithm (version 0.10.0)(109). Marker genes were assigned to each cluster using the FindAllMarkers function from Seurat. Statistical significance was inferred using the Wilcoxon test after FDR adjustment. For the purposes of our analyzes, cortical cell clusters expressing known markers of each cortical zone (zG, zF, and zR in humans and zG, zF and x-Zone in mice) were collapsed together and marker genes were recalculated (Supplemental Figure 5a-b).

### Protein expression and purification

WNT carriers, receptors (FZD cysteine-rich domains), coreceptors (GPC ectodomain and ROR extracellular domain), and WNT-carrier complexes were stably expressed in HEK293 cells and were purified from conditioned media as previously described(65). Expression of the purified proteins were confirmed by SDS-PAGE, Coomassie staining or immunoblotting using rabbit monoclonal or polyclonal antibodies against WNT5A/B (Cell Signaling, #2530S) or WNT2B (Abcam, #ab203225).

### WNT release and canonical WNT activity assays

HEK293 cells stably expressing WNT or NanoLuc (NL) luciferase tagged WNT were washed thrice with serum-free DMEM and were incubated at the indicated time points with the indicated WNT carriers and GPC ectodomains. Conditioned media was collected in duplicate at each time point, centrifuged to remove cellular debris, and subjected to Nano-Glo Luciferase Assay Substrate (Promega), according to the manufacturer’s instructions. NL-tagged WNT released into the media was normalized using the total NL signal in the corresponding cells, harvested at the end of the time course.

Canonical activity of WNT conditioned media, purified WNT-carrier complexes or R-spodin3 (R&D Systems 4120-RS), was measured after 24h incubation in MEF cells stably expressing the TopFlash reporter system, which consists of a firefly luciferase under the control of a TCF response element and Renilla luciferase expressed constitutively(65, 66). Luminescence was assessed in cell lysates in duplicate by Dual-Glo Luciferase Assay System (Promega), using a Victor3 Multilabel plate reader (Perkin-Elmer). Wnt pathway activation was calculated as the ratio of firefly to Renilla luminescence, normalized to untreated cells (serum-free DMEM), with error bars representing SD.

### WNT transfer on beads

Conditioned media from NL-WNT2B-expressing cells were collected after 48h transfection with plasmids encoding HT7-tagged carriers or entire GPC ectodomains and were captured on HaloLink beads (Promega) as previously described(65). NL-WNT2B-carrier beads (5µL) were incubated with 5μM of purified FZD-CRDs, diluted in HBS (20mM HEPES, pH 7.5; 150mM NaCl) and preincubated with a 20-fold excess of HaloLink-amine to block the HaloTag7 (HT7)(110). Beads and FZD-CRDs supernatant were then tumbled at room temperature for 2, 5, 15 and 30min timepoint. At the end of the time course, NL luminescence in the supernatant aliquots and on beads was measured as described above (WNT release assay). NL-WNT released in the supernatant was represented as percentage of the total NL signal on beads.

### Immunoprecipitation

Purified FLAG-tagged ROR1-ECD and ROR2-ECD coreceptors (2.5μM) were incubated for 3h at room temperature with purified WNT-carrier complexes or carriers alone (5μM), diluted in TBS with 2mM CaCl_2_ and 0.2% DDM. After 3h incubation, the samples were tumbled overnight at 4°C and immunoprecipitated on anti-FLAG beads. After washing the beads three times with 2mM CaCl_2_ and 0.2% DDM, bound proteins were eluted in elution buffer (20mM HEPES, pH 7.5; 200mM NaCl; 5mM EDTA; 100µg/mL FLAG or HPC peptide) and were analyzed by SDS-PAGE followed by immunoblotting using rabbit monoclonal or polyclonal antibodies against WNT3A (Cell Signaling, #2721S), WNT5A/B (Cell Signaling, #2530S) or, WNT2B (Abcam, #ab203225), and anti-mouse monoclonals against FLAG-M1 and anti-HPC, a generous gift from Andrew C Kruse (Harvard Medical School). RhoA/Rac1 activity assay. Wild-type (WT), FZD(1–10)^KO^(79), or ROR(1–2)^KO^(65). HEK293 cells were pretreated overnight with the Porcupine (PORCN) inhibitor, IWP-2 (2μM, Sigma), and/or transfected for 24h with the indicated FZD receptor and ROR coreceptor, followed by 3h incubation with 2μM of purified GPC4 alone or in complex with WNT3A, WNT5A or WNT2B, diluted in serum-free DMEM. Cells were then lysed with 1x assay/lysis buffer (Cell Biolabs, #STA-405) and clarified by centrifugation at 14,000×*g* for 10 min at 4 °C. Activity of RhoA and Rac1 was assayed using RhoA/Rac1/Cdc42 Activation Assay Combo Kit (Cell Biolabs, #STA-405). Briefly, clarified lysates were incubated with Rhotekin RBD beads or PAK1 PBD beads for 24h with gentle agitation at 4 °C. Beads were then washed three times with 1x assay/lysis buffer and Rhotekin RBD/GTP-RhoA or PAK PBD/GTP-Rac1 was analyzed by SDS-PAGE followed by immunoblotting using specific anti-mouse Rac1 or RhoA antibodies, according to the manufacturer’s protocol.

### Assessment of RhoA activation by Dual-Glo Luciferase reporter gene assay

RhoA activity was assessed using the Serum Response Factor Response Element (SRF-RE) pGL4.34 (Promega, E1350), which drives the transcription of the luciferase reporter gene *luc2P* in response to activation of Serum Response Factor that triggers RhoA GTPase. The pGL4.34 firefly luciferase reporter (SRF-RE) and the renilla luciferase thymidine kinase (pRL-TK) reporter (Promega, E2261) were co-transfected into HEK-293 (8×10^3^ cells/well) cells cultured in a 96-well plate for 24h and 48h, respectively, as previously described(111). Following transfection, the cells were treated with WNT-carrier complexes for an additional 6 hours of incubation. Luciferase activity was measured using the Dual-Luciferase Reporter Assay System (Promega), and the ratio of firefly luciferase activity to Renilla luciferase activity was calculated for each well. Experiments were performed in duplicate and repeated three times.

### Human subjects

The study protocols were approved by the Boston Children’s Institutional Review Board (P10-02-0053 and P00020529). Written informed consent was obtained from the guardians of all pediatric participants prior to their inclusion in the study. Guardians were provided with detailed information regarding the study objectives, procedures, potential risks, and benefits. No compensation was provided for participation in this study.

### Genome-Wide Association Stud

A case-control multi-ancestry cohort analysis was performed using the *All of Us* Database. We identified people who had hyperaldosteronism by performing a keyword search on the conditions listed by *All of Us*, as well as in the ICD10 codes for the participants with the keywords: ‘hyperaldosteronism,’ and ‘resistant hypertension’. Of these, individuals with short read whole genome sequence data were selected as cases. If samples were from related individuals (kinship score > 0.1), we selected only one member from the family to form a set of unrelated individuals. With these criteria, and specific exclusion criteria **(Appendix 1**), we identified 271 cases.

For the control cohort, we sought to select individuals without primary hyperaldosteronism (PA). In addition, we excluded individuals with hypertension since PA is under-diagnosed and is often diagnosed as hypertension without a specific etiology. From the 413,457 participants in the *All of Us* study, 245,195 participants have short-read whole genome data. After excluding cases, we excluded participants who had an ICD 10 code indicating the existence of hypertension of any kind. We also excluded participants whose EHR records reflected elevated BP measures and participants who had at least one elevated BP measure (systolic blood pressure >= 140 or diastolic blood pressure >= 90) in their recorded Labs & Measurements. With these criteria, a total of 74,354 controls were identified (**Fig. A1**).

Sixteen principal components (PC), precalculated and provided with *All of Us* genetic data, were used to infer ancestry and to control for population stratification during association testing. Rather than splitting the small number of cases further by the genetically-inferred population labels provided by *All of Us*, we included all cases in a single multi-ancestry analysis, selecting five controls for each case that had the lowest Mahalanobis distance to the case in the PC space. The distribution of ancestry and sex between cases and matched controls is presented in **Appendix 2** and **Table A1**. The genotype distribution between cases and matched controls is presented in **Table A2**. We then tested the association in this case-control cohort between rs3790604 and PA by using a logistic regression model, with the 16 PCs as covariates. The input genotype data set we used includes variants that are frequent in the computed ancestry subpopulations (population-specific allele frequency > 1% OR population-specific allele count > 100; known as the ACAF callset).

## APPENDIX 1: Inclusion / Exclusion Criteria for Cases & Controls

INCLUDE Individuals from *All of Us* with the following:

Conditions:

1. Hyperaldosteronism
2. Secondary hyperaldosteronism
3. Primary aldosteronism
4. Resistant hypertensive disorder

ICD 10 Codes:

1. E26.0 (Primary hyperaldosteronism)
2. E26.1 (Secondary hyperaldosteronism)
3. E26.89 (Other hyperaldosteronism)
4. E26.9 (Hyperaldosteronism, unspecified)
5. I15.2 (Hypertension secondary to endocrine disorders)
6. I15.8 (Other secondary hypertension)
7. I15.9 (Secondary hypertension, unspecified)

From the resulting set of cases, EXCLUDE samples that could be classified as cases or controls, but the documented evidence does not categorize them as one or the other. The specific phrases and conditions that were excluded:

1. Eclampsia with pre-existing hypertension
2. Pre-existing hypertensive heart & chronic kidney disorder
3. Secondary hypertension
4. Hypertension secondary to endocrine disorder
5. Benign secondary hypertension
6. Benign secondary renovascular hypertension
7. Malignant secondary renovascular hypertension
8. Malignant secondary hypertension

Pre-existing secondary hypertension complicating pregnancy, childbirth and puerperium The inclusion / exclusion criteria are shown in **Figure A1**.

**Figure A1:**
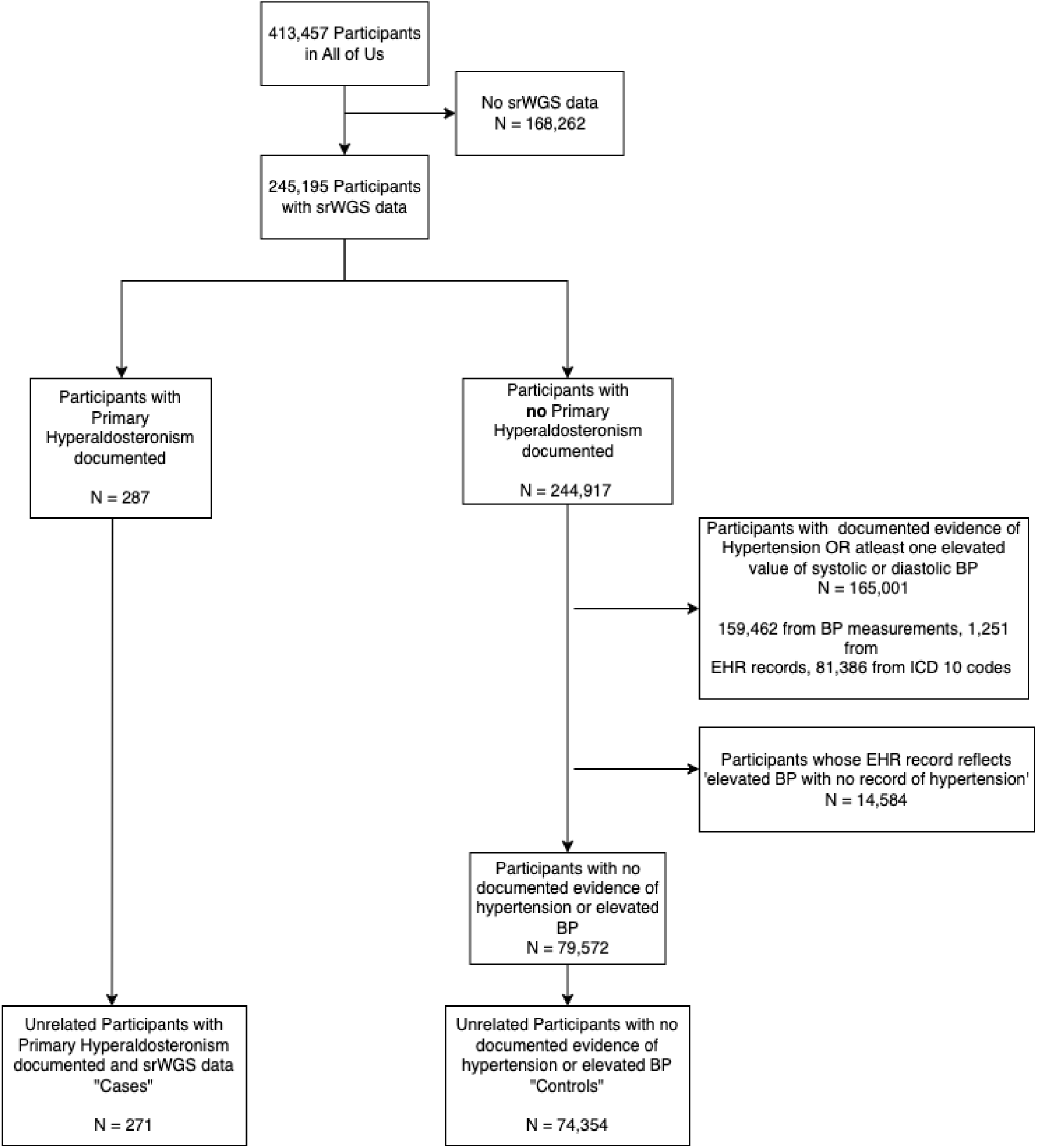
Inclusion / Exclusion Criteria for the Study.

## APPENDIX 2: Ancestry & Sex-Distribution

**Table A1:**
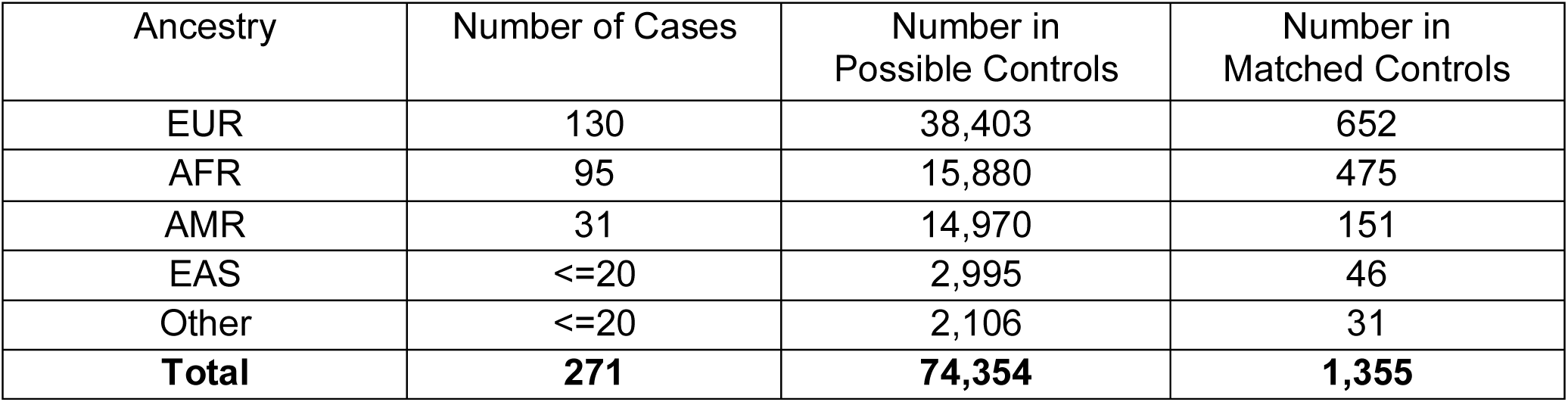
Ancestry Distribution within Case & Control Cohorts.

| Ancestry | Number of Cases | Number in Possible Controls | Number in Matched Controls |
| --- | --- | --- | --- |
| EUR | 130 | 38,403 | 652 |
| AFR | 95 | 15,880 | 475 |
| AMR | 31 | 14,970 | 151 |
| EAS | <=20 | 2,995 | 46 |
| Other | <=20 | 2,106 | 31 |
| **Total** | **271** | **74,354** | **1,355** |

The distribution of samples shown above is from the unrelated set of cases, the unrelated set of controls that could be selected, and the unrelated set of controls that were selected based on their distance from the cases in PC-space.

Within the case cohort, we identified 149 females (54.98%), and 116 (42.80%) males. The distribution of sex is approximately equal for the EUR ancestry (47.97% of the case cohort, with 65 (50%) females and 60 (46.15%) males), but predominantly female in those of AFR ancestry (35.06% of the case cohort, with 59 (62.10%) females and 35 (36.84%) males).

Within the possible controls, this trend reverses: the EUR ancestry had an increased proportion of females (51.65% of the possible controls, with 25,108 (65.38%) females and 12,585 (32.77%) males), while there was a balanced sex-distribution amongst the AFR ancestry subset (21.36% of the possible controls, with 7,779 (51.96%) females and 6,859 (45.82%)). We also observe an increased proportion of females (46,971 females (63.20%), 26,002 males (34.98%)).

Among the matched controls, the sex-distribution was similar to that of the set of possible controls (844 (62.33%) female, 484 (35.74%) male) and the trend of sex-distribution of the possible control set remained reflected in the EUR ancestry (441 (67.64%) female, 201 (30.83%) male) and AFR ancestry (246 (51.79%) females, 217 (45.68%) males) of the matched controls.

Importantly, the distribution of ancestry of the matched controls was similar to the matched cases (EUR ancestry: 48.12% of the matched control cohort, AFR ancestry: 35.05% of the matched control cohort) – a result of and a basic indicator of our ancestry-based matching.

**Table A2:** Genotype Distribution within Case & Control Cohorts. The distribution for SNP rs3790604 (GRCh38:1:112,504,257:C=>A, GRCh37:1:113,046,879:C=>A) is as follows:

| Rs3790604 genotypes | C/C | C/A | A/A | None |
| --- | --- | --- | --- | --- |
| Cases | 225 | 42 | 1 | 3 |
| Controls | 1194 | 141 | 6 | 14 |

---

**Supplemental Figure 1.**
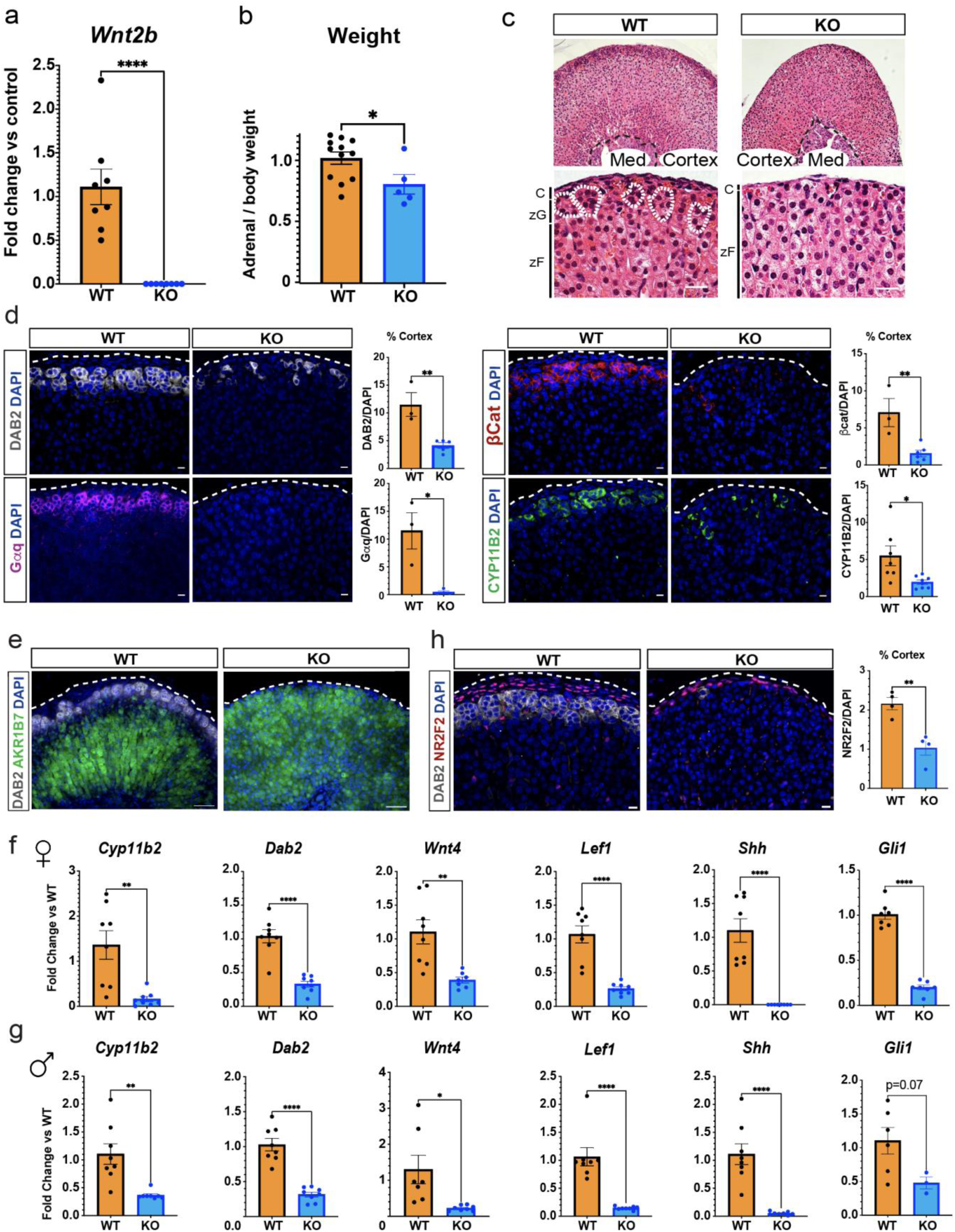
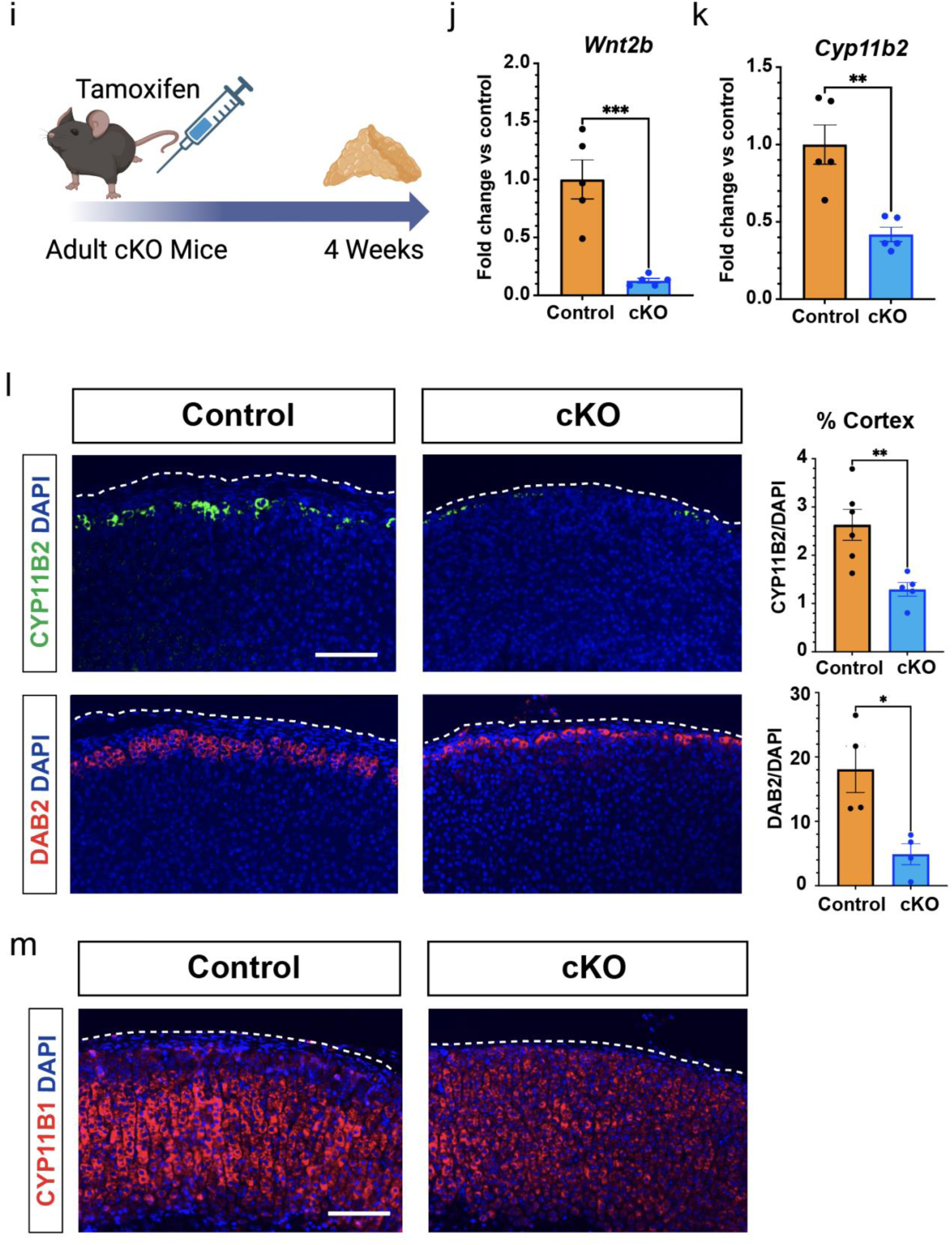
WNT2B deficiency results in a dysmorphic zG in mice. a. QRT-PCR was performed on WT and KO male adrenals (n=8 WT, n=8 KO). Two-tailed Student’s t-test. ****p < 0.0001. Data are represented as mean ± SEM. b. Adrenal weight normalized to body weight from male mice (n=12 WT, n=5 KO). Two -tailed Student’s t-test. *p < 0.05. Data are represented as mean fold change ± SEM. c. Representative H&E images of WT and KO male adrenals. Scale bar: 10μm. C, capsule; zG, zona glomerulosa; zF, zona fasciculata; Med, medulla. d. Representative images and quantification from male adrenals stained for DAB2 (gray, n=3 WT, n=5 KO), Gαq (magenta, n=3 WT, n=4 KO), β-catenin (β-cat, red, n=3 WT, n=6 KO) and CYP11B2 (green, n=7 WT, n=7 *Wnt2b* KO). Positive cells were quantified and normalized to nuclei (DAPI) in the cortex. Scale bars: 10μm. Two-tailed Student’s t-test. *p < 0.05; **p < 0.01. Data are represented as mean ± SEM. e. Representative images stained for DAB2 (gray), AKR1B7 (green) and DAPI (blue) from WT and KO adrenals. Scale bar: 50μm f. QRT-PCR was performed on WT and KO female adrenals for *Cyp11b2* (n=8 WT, n=7 KO), *Dab2* (n=8 WT, n=7 KO), *Wnt4* (n=8 WT, n=7 KO), *Lef1* (n=8 WT, n=8 KO), *Shh* (n=8 WT, n=8 KO) and *Gli1* (n=7 WT, n=7 KO). Two-tailed Student’s t-test. **p < 0.01; ****p < 0.0001. Data are represented as mean ± SEM. g. QRT-PCR was performed on WT and KO male adrenals for *Cyp11b2* (n=8 WT, n=8 KO), *Dab2* (n=8 WT, n=8 KO), *Wnt4* (n=7 WT, n=8 KO), *Lef1* (n=8 WT, n=8 KO), *Shh* (n=8 WT, n=8 KO) and *Gli1* (n=6 WT, n=3 KO). Two-tailed Student’s t-test. *p < 0.05; **p < 0.01; ****p < 0.0001. Data are represented as mean ± SEM. h. Representative images and quantification of immunostaining for DAB2 (gray) and NR2F2 (red) from WT and KO adrenals (n=4 WT, n=4 KO). Positive cells were quantified and normalized to nuclei (DAPI, blue) in the cortex. Scale bars: 10μm. Two-tailed Student’s t-test. **p < 0.01. Data are represented as mean ± SEM. i. Treatment protocol of adult cKO mice at 6-7 weeks of age with tamoxifen and adrenal harvest after 4 weeks. j. QRT-PCR was performed for *Wnt2b* in adrenals (n=5 Control and n=5 cKO) 4 weeks following tamoxifen injection. Two-tailed Student’s t-test. ***p < 0.001. Data are represented as mean ± SEM. k. QRT-PCR was performed for *Cyp11b2* in adrenals (n=5 Control and n=5 cKO) 4 weeks following tamoxifen injection. Two-tailed Student’s t-test. **p < 0.01. Data are represented as mean ± SEM. l. Representative images and quantification from adrenals stained for CYP11B2 (green, n=6 Control, n=5 cKO and DAB2 (red, n=4 Control, n=4 cKO) 4 weeks following tamoxifen injection. Positive cells were quantified and normalized to nuclei (DAPI) in the cortex. Scale bar: 100μm. Two-tailed Student’s t-test. *p < 0.05; **p < 0.01. Data are represented as mean ± SEM. m. Representative images from Control and cKO adrenals stained for CYP11B1 (red) and DAPI (blue) 4 weeks following tamoxifen injection. Scale bar: 100μm.

**Supplemental Figure 2.**
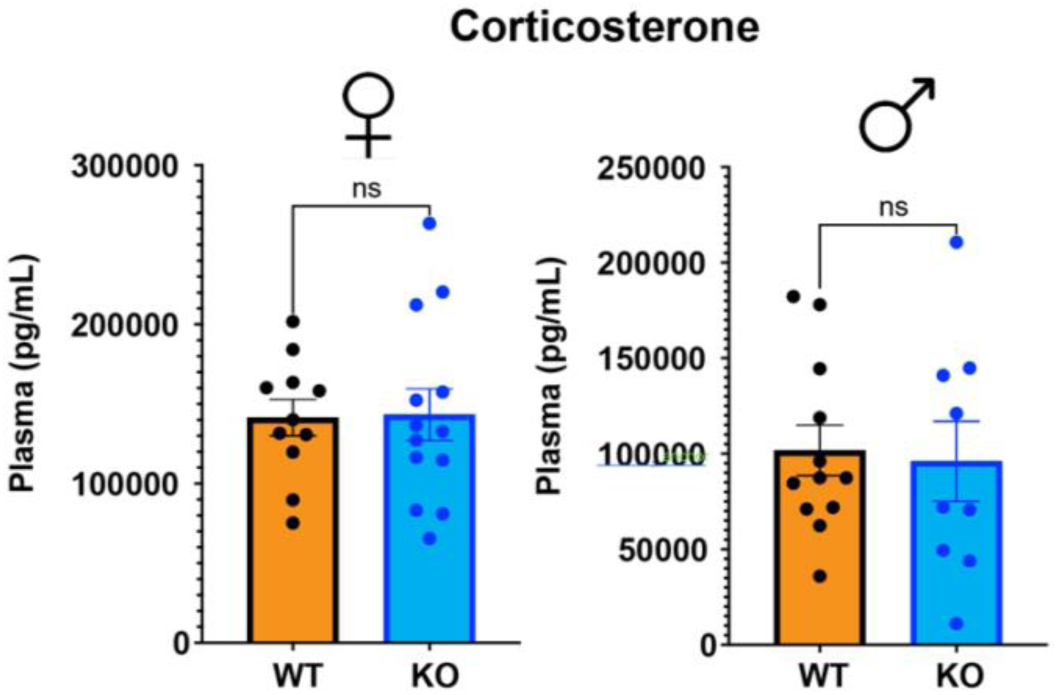
WNT2B deficiency does not affect corticosterone levels in mice. Plasma corticosterone levels (female, n=11 WT, n=13 KO; male, n=12 WT, n=9 KO). Two-tailed Student’s t-test. ns, not significant. Data are represented as mean ± SEM.

**Supplemental Figure 3.**
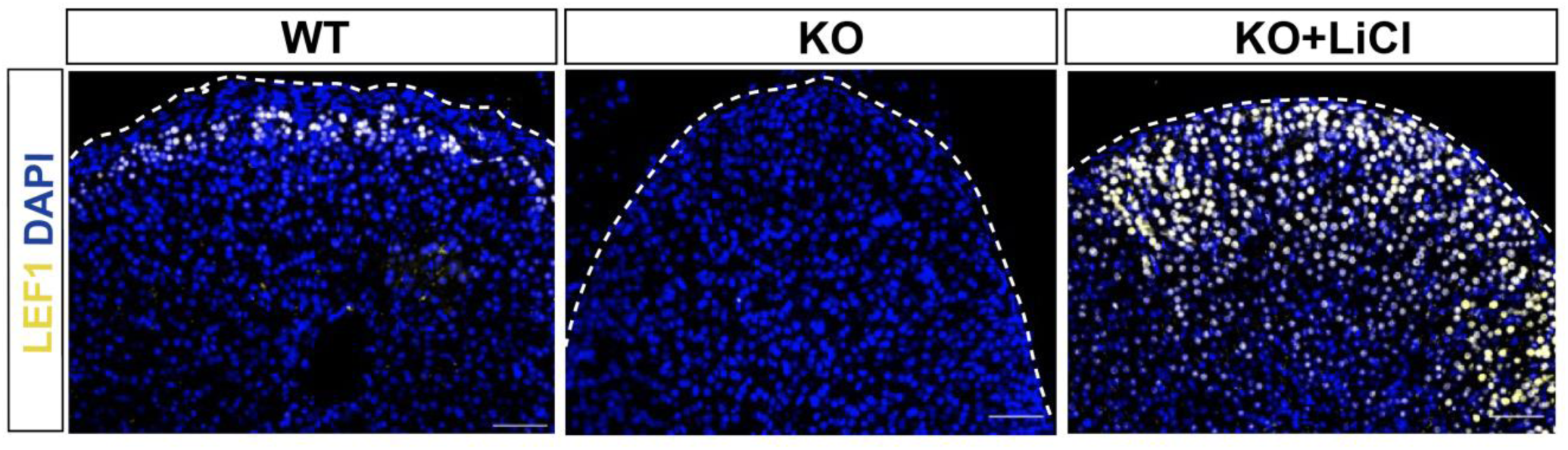
Representative images from female adrenals immunostained for LEF1 (yellow) and DAPI (blue) from WT, KO and KO+LiCL mice. Scale bar: 50μm

**Supplemental Figure 4.**
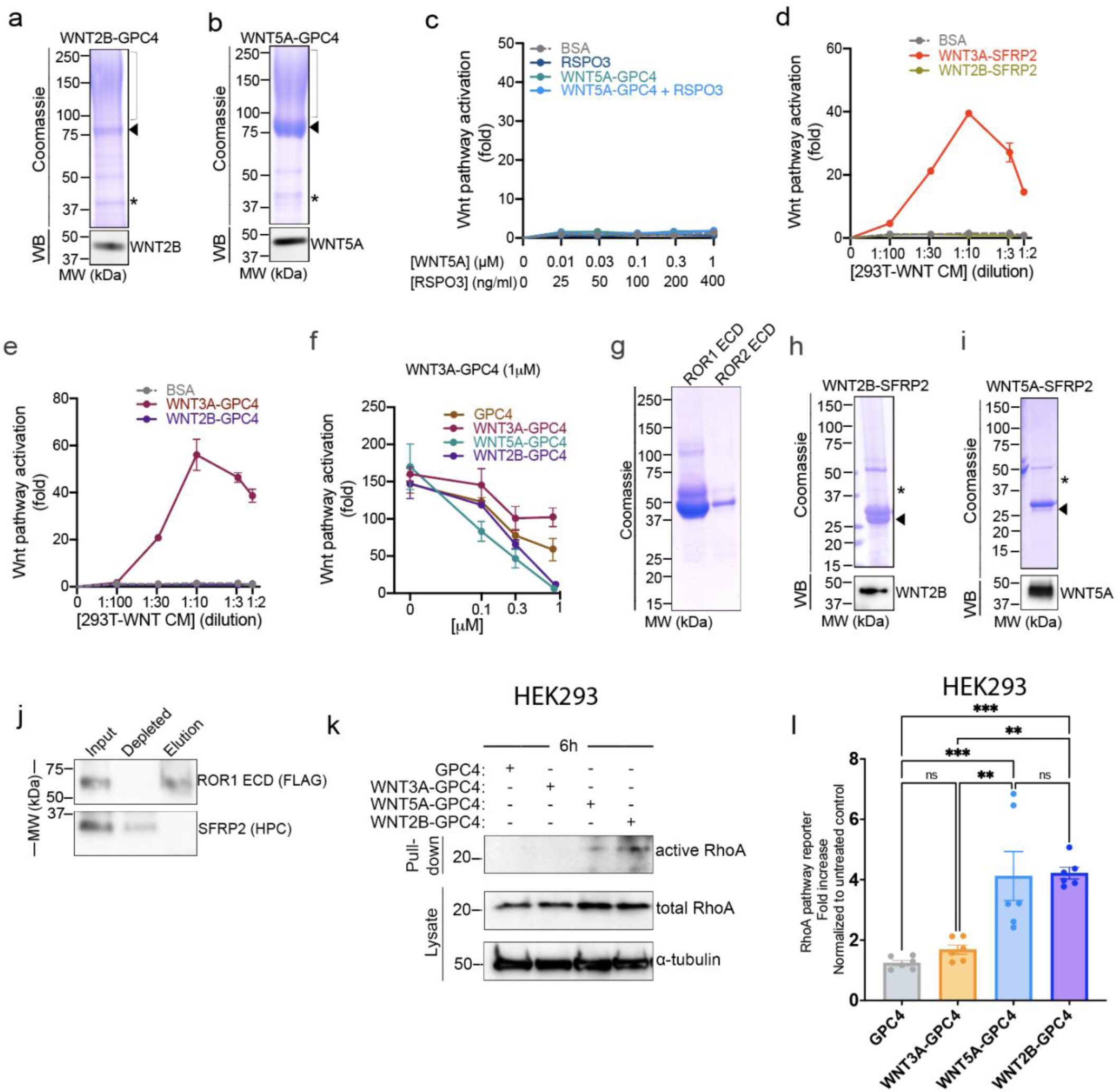
Characterization of WNT2B as a non-canonical ligand. a. WNT2B-GPC4 ectodomain, C-terminally tagged with HaloTag7 (HT7) and HPC tag, was affinity purified from conditioned media on an anti-HPC antibody matrix, and analyzed by SDS-PAGE, followed by Coomassie staining or anti-WNT2B immunoblotting (WB). Arrowhead indicates unmodified GPC4, bracket indicates glycosaminoglycan (GAG)-modified species, and asterisks indicate WNT2B protein. b. As in (a), but with WNT5A in complex with GPC4, and with anti-WNT5A immunoblotting. c. R-Spondin 3 (RSPO3, 0, 25, 100, 200 and 400ng/ml) or purified WNT5A-GPC4 complex (0.01, 0.03, 0.1, 0.3 and 1μM with respect to WNT3A) with or without RSPO3 (400ng/ml) was added to Wnt reporter cells. After 24h, Wnt pathway activity was measured by luciferase assay. Incubation with BSA served as negative control. WNT5A-GPC4 does not activate canonical Wnt signaling, even when incubated with RSPO3. Points represent average activation for two biological replicates, normalized to untreated cells, and error bars represent SD. d. SFRP2 (1μM) was added in serum-free media in WNT3A- or WNT2B-expressing HEK293 cells. Serial dilutions of the conditioned media were then added to Wnt reporter cells, and Wnt pathway activity was measured by Dual-Glo luciferase 24h later. BSA (1μM) served as negative control. WNT2B released by SFRP2 is unable to activate canonical Wnt signaling, in contrast to WNT3A-SFRP2 conditioned media. Points represent average activation for two biological replicates, normalized to the negative control, and error bars represent SD. e. As in (d), but WNT-expressing cells were incubated with 1μM of GPC4. f. As in (Fig. 4d), but purified WNT3A-GPC4 complex (1μM) was mixed with the indicated concentrations of GPC4 alone or in complex with WNT3A, WNT5A or WNT2B. WNT3A-SFRP2 activity is abolished by WNT5A-GPC4 and WNT2B-GPC4 complexes in a dose-dependent manner, which contrasts GPC4 alone or WNT3A-GPC4 complex. g. Extracellular domains (ECD) of ROR1 and ROR2, N-terminally tagged with a FLAG tag, were affinity purified from conditioned media on an anti-FLAG antibody matrix. Purified proteins were analyzed by SDS-PAGE and Coomassie staining. h. As in (a), but with WNT2B in complex with SFRP2, C-terminally tagged with 8x-His tag and HPC taI. i. As in (b), but with WNT5A in complex with SFRP2. j. Purified SFRP2 (5μM) was incubated with FLAG-tagged ROR1-ECD (2.5μM), followed by immunoprecipitation with antibodies against the FLAG tag. Samples were analyzed by SDS-PAGE and immunoblotting. SFRP2 does not interact with ROR1-EcD. k. Activity of RhoA in cell lysates of HEK293 cells treated for 6h with GPC4 alone or in complex with WNT3A, WNT5A or WNT2B (2μM) was assessed by Rhotekin-RBD pull-down assay. RhoA endogenous levels are shown in the lysates. Both WNT5A-GPC4 and WNT2B-GPC4 complexes induce activity of RhoA, in contrast to GPC4 alone or in complex with WNT3A. Blotting for α-tubulin served as loading control. l. HEK293 cells were co-transfected with the firefly luciferase reporter (pGL4.34) and the renilla luciferase thymidine kinase reporter (pRL-TK). They were then used to assay RhoA activation by purified GPC4 alone or in complex with WNT3A, WNT5A, and WNT2B (1 μM). We found that the activity of RhoA is induced by WNT2B-GPC4 or WNT5A-GPC complexes, but not by WNT3A-GPC4 or GPC4 alone. The bars represent the average from three independent experiments performed in duplicate, normalized to untreated cells. Statistical significance was determined using one-way ANOVA with Tukey’s post-test (ns, not significant; **p < 0.01; ***p < 0.001). Data are represented as mean ± SEM.

**Supplemental Figure 5.**
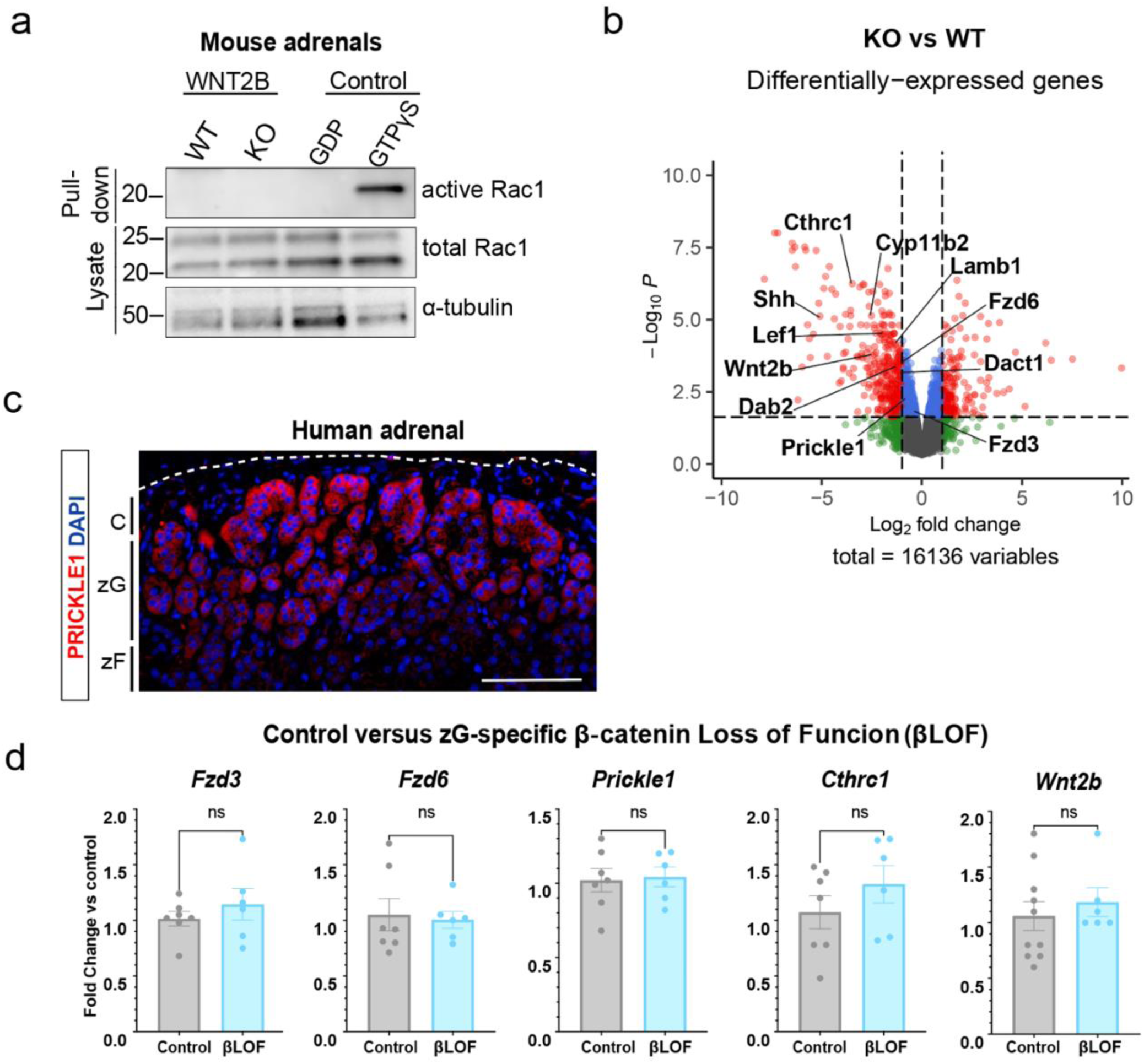
WNT2B deficiency disrupts Wnt/PCP signaling in the adrenal. a. Activity of Rac1 in WT and KO adrenals assessed by Rhotekin-RBD pull-down assay using adrenal lysates. GTPγS and GDP treated adrenal lysates served as positive and negative controls, respectively. Total Rac1 and α-tubulin served as loading controls. b. Volcano plot showing differentially-expressed genes between WT and KO adrenals. Dots representing genes down- and up-regulated in KO are displayed on the left and right sides of the plot, respectively. Red dots represent genes that exhibit a fold-change > 2-fold with a FDR-adjusted p-value < 0.05. Selected zonal markers, including zG genes, are indicated. c. Representative image stained for PRICKLE1 (red) and DAPI (blue) from human adrenals. Scale bar: 100μm. C, capsule; zG, zona glomerulosa; zF, zona fasciculata. d. QRT-PCR was performed in WT and zG-specific β−catenin LOF adrenals for Fzd3 (n=7 Control, n=6 βLOF), Fzd6 (n=7 Control, n=6 βLOF), Prickel1 (n=7 Control, n=6 βLOF), Cthrc1 (n=7 Control, n=6 βLOF), and Wnt2b (n=10 Control, n=6 βLOF). Two-tailed Student’s t-test. ns, not significant. Data are represented as mean ± SEM.

**Supplemental Figure 6.**
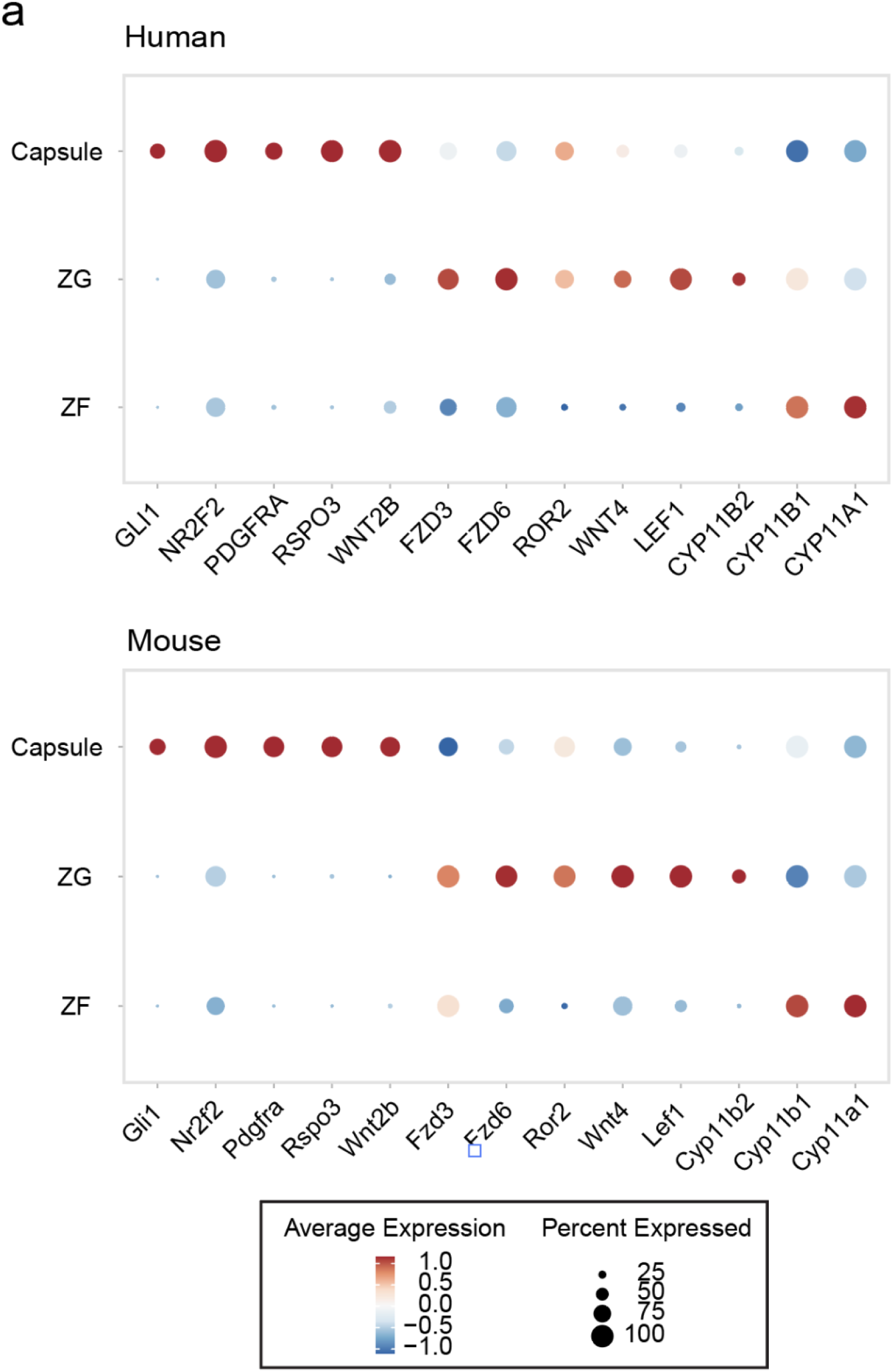
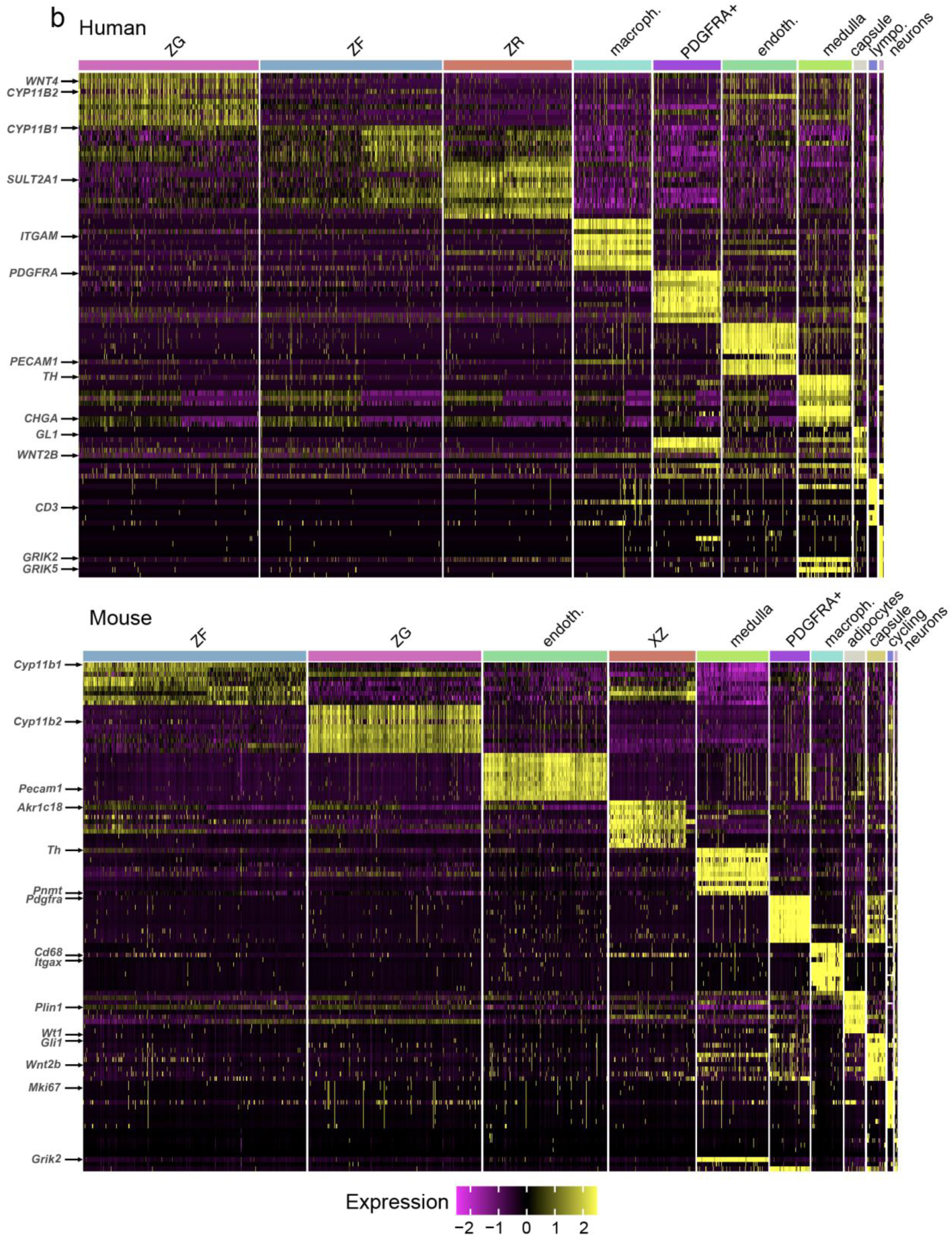
a. Dot plot showing average expression of genes in the capsule, zG or zF from human or mouse adrenals. b. Heatmap visualization showing gene expression patterns of cellular clusters identified in human and mouse adrenals.

## References

1. Hammer GD, Basham KJ. Stem cell function and plasticity in the normal physiology of the adrenal cortex. Mol Cell Endocrinol. 2021;519:111043.

2. Zhu Y, Zhang X, Hu C. Structure of rosettes in the zona glomerulosa of human adrenal cortex. J Anat. 2023;243(4):684–689.

3. Leng S, et al. β-Catenin and FGFR2 regulate postnatal rosette-based adrenocortical morphogenesis. Nat Commun. 2020;11(1):1680.

4. Zelander T. The ultrastructure of the adrenal cortex of the mouse. Zeitschrift für Zellforschung und Mikroskopische Anatomie. 1957;46(6):710–716.

5. Freedman BD, et al. Adrenocortical Zonation Results from Lineage Conversion of Differentiated Zona Glomerulosa Cells. Dev Cell. 2013;26(6):666–673.

6. White PC. Aldosterone synthase deficiency and related disorders. Mol Cell Endocrinol. 2004;217(1– 2):81–87.

7. Kayes-Wandover KM, et al. Congenital Hyperreninemic Hypoaldosteronism Unlinked to the Aldosterone Synthase (CYP11B2) Gene. J Clin Endocrinol Metab. 2001;86(11):5379–5382.

8. Lucas C, et al. Loss of LGR4/GPR48 causes severe neonatal salt wasting due to disrupted WNT signaling altering adrenal zonation. Journal of Clinical Investigation. 2023;133(4). 10.1172/JCI164915.

9. Hui E, et al. The clinical significance of aldosterone synthase deficiency: report of a novel mutation in the CYP11B2 gene. BMC Endocr Disord. 2014;14(1):29.

10. Wu V-C, et al. The prevalence of CTNNB1 mutations in primary aldosteronism and consequences for clinical outcomes. Sci Rep. 2017;7(1):39121.

11. Azizan EAB, Drake WM, Brown MJ. Primary aldosteronism: molecular medicine meets public health. Nat Rev Nephrol. 2023;19:788–806.

12. Ruiz-Sánchez JG, et al. Clinical manifestations and associated factors in acquired hypoaldosteronism in endocrinological practice. Front Endocrinol (Lausanne). 2022;13. 10.3389/fendo.2022.990148.

13. Vaidya A, et al. Primary Aldosteronism: State-of-the-Art Review. Am J Hypertens. 2022;35(12):967– 988.

14. Scholl UI. Genetics of Primary Aldosteronism. Hypertension. 2022;79(5):887–897.

15. Nanba K, et al. Molecular Heterogeneity in Aldosterone-Producing Adenomas. J Clin Endocrinol Metab. 2016;101(3):999–1007.

16. Boulkroun S, et al. Adrenal Cortex Remodeling and Functional Zona Glomerulosa Hyperplasia in Primary Aldosteronism. Hypertension. 2010;56(5):885–892.

17. Dekkers T, et al. Adrenal Nodularity and Somatic Mutations in Primary Aldosteronism: One Node Is the Culprit? J Clin Endocrinol Metab. 2014;99(7):E1341–E1351.

18. Iacobone M, et al. Unilateral adrenal hyperplasia: A novel cause of surgically correctable primary hyperaldosteronism. Surgery. 2012;152(6):1248–1255.

19. Seccia TM, et al. The Biology of Normal Zona Glomerulosa And Aldosterone-Producing Adenoma: Pathological Implications. Endocr Rev. 2018;39(6):1029–1056.

20. Steinhart Z, Angers S. Wnt signaling in development and tissue homeostasis. Development. 2018;145(11). 10.1242/dev.146589.

21. Yu M, et al. The evolving roles of Wnt signaling in stem cell proliferation and differentiation, the development of human diseases, and therapeutic opportunities. Genes Dis. 2024;11(3):101026.

22. Sugimura R, et al. Noncanonical Wnt Signaling Maintains Hematopoietic Stem Cells in the Niche. Cell. 2012;150(2):351–365.

23. Wiese KE, Nusse R, van Amerongen R. Wnt signalling: conquering complexity. Development. 2018;145(12). 10.1242/dev.165902.

24. Janda CY, et al. Structural Basis of Wnt Recognition by Frizzled. Science (1979). 2012;337(6090):59–64.

25. Cheng Z, et al. Crystal structures of the extracellular domain of LRP6 and its complex with DKK1. Nat Struct Mol Biol. 2011;18(11):1204–1210.

26. Cong F, Schweizer L, Varmus H. Wnt signals across the plasma membrane to activate the β-catenin pathway by forming oligomers containing its receptors, Frizzled and LRP. Development. 2004;131(20):5103–5115.

27. Janda CY, et al. Surrogate Wnt agonists that phenocopy canonical Wnt and β-catenin signalling. Nature. 2017;545(7653):234–237.

28. Sheetz JB, et al. Structural Insights into Pseudokinase Domains of Receptor Tyrosine Kinases. Mol Cell. 2020;79(3):390–405.e7.

29. Ho H-YH, et al. Wnt5a–Ror–Dishevelled signaling constitutes a core developmental pathway that controls tissue morphogenesis. Proceedings of the National Academy of Sciences. 2012;109(11):4044– 4051.

30. Martinez S, et al. The PTK7 and ROR2 Protein Receptors Interact in the Vertebrate WNT/Planar Cell Polarity (PCP) Pathway. Journal of Biological Chemistry. 2015;290(51):30562–30572.

31. Berthon A, et al. Constitutive β-catenin activation induces adrenal hyperplasia and promotes adrenal cancer development. Hum Mol Genet. 2010;19(8):1561–1576.

32. Basham KJ, et al. A ZNRF3-dependent Wnt/β-catenin signaling gradient is required for adrenal homeostasis. Genes Dev. 2019;33(3–4):209–220.

33. Finco I, Lerario AM, Hammer GD. Sonic Hedgehog and WNT Signaling Promote Adrenal Gland Regeneration in Male Mice. Endocrinology. 2018;159(2):579–596.

34. Pignatti E, et al. Beta-Catenin Causes Adrenal Hyperplasia by Blocking Zonal Transdifferentiation. Cell Rep. 2020;31(3):107524.

35. Heikkilä M, et al. Wnt-4 Deficiency Alters Mouse Adrenal Cortex Function, Reducing Aldosterone Production. Endocrinology. 2002;143(11):4358–4365.

36. Kim AC, et al. Targeted disruption of β-catenin in Sf1-expressing cells impairs development and maintenance of the adrenal cortex. Development. 2008;135(15):2593–2602.

37. Drelon C, et al. PKA inhibits WNT signalling in adrenal cortex zonation and prevents malignant tumour development. Nat Commun. 2016;7(1):12751.

38. Heaton JH, et al. Progression to Adrenocortical Tumorigenesis in Mice and Humans through Insulin-Like Growth Factor 2 and β-Catenin. Am J Pathol. 2012;181(3):1017–1033.

39. Borges KS, et al. Wnt/β-catenin activation cooperates with loss of p53 to cause adrenocortical carcinoma in mice. Oncogene. 2020;39(30):5282–5291.

40. Lyraki R, et al. Crosstalk between androgen receptor and WNT/β-catenin signaling causes sex-specific adrenocortical hyperplasia in mice. Dis Model Mech. 2023;16(6). 10.1242/dmm.050053.

41. Vidal V, et al. The adrenal capsule is a signaling center controlling cell renewal and zonation through *Rspo3*. Genes Dev. 2016;30(12):1389–1394.

42. Zhou J, et al. Somatic mutations of GNA11 and GNAQ in CTNNB1-mutant aldosterone-producing adenomas presenting in puberty, pregnancy or menopause. Nat Genet. 2021;53(9):1360–1372.

43. Nanba K, et al. Double somatic mutations in CTNNB1 and GNA11 in an aldosterone-producing adenoma. Front Endocrinol (Lausanne). 2024;15. 10.3389/fendo.2024.1286297.

44. Teo AED, et al. Pregnancy, Primary Aldosteronism, and Adrenal CTNNB1 Mutations. New England Journal of Medicine. 2015;373(15):1429–1436.

45. Scholl UI, et al. Novel somatic mutations in primary hyperaldosteronism are related to the clinical, radiological and pathological phenotype. Clin Endocrinol (Oxf). 2015;83(6):779–789.

46. Naito T, et al. Genetic Risk of Primary Aldosteronism and Its Contribution to Hypertension: A Cross-Ancestry Meta-Analysis of Genome-Wide Association Studies. Circulation. 2023;147(14):1097–1109.

47. Inoue K, et al. Primary Aldosteronism and Risk of Cardiovascular Outcomes: Genome-Wide Association and Mendelian Randomization Study. J Am Heart Assoc. 2024;13(15). 10.1161/JAHA.123.034180.

48. Bick AG, et al. Genomic data in the All of Us Research Program. Nature. 2024;627(8003):340–346.

49. The “All of Us” Research Program. New England Journal of Medicine. 2019;381(7):668–676.

50. Lin Y, et al. Induction of ureter branching as a response to Wnt-2b signaling during early kidney organogenesis. Developmental Dynamics. 2001;222(1):26–39.

51. Drelon C, et al. Adrenal cortex tissue homeostasis and zonation: A WNT perspective. Mol Cell Endocrinol. 2015;408:156–164.

52. Tsukiyama T, Yamaguchi TP. Mice lacking Wnt2b are viable and display a postnatal olfactory bulb phenotype. Neurosci Lett. 2012;512(1):48–52.

53. Romero DG, et al. Disabled-2 Is Expressed in Adrenal Zona Glomerulosa and Is Involved in Aldosterone Secretion. Endocrinology. 2007;148(6):2644–2652.

54. Aigueperse C, et al. Cyclic AMP regulates expression of the gene coding for a mouse vas deferens protein related to the aldo-keto reductase superfamily in human and murine adrenocortical cells. Journal of Endocrinology. 1999;160(1):147–154.

55. King P, Paul A, Laufer E. Shh signaling regulates adrenocortical development and identifies progenitors of steroidogenic lineages. Proceedings of the National Academy of Sciences. 2009;106(50):21185–21190.

56. Huang C-CJ, et al. Progenitor Cell Expansion and Organ Size of Mouse Adrenal Is Regulated by Sonic Hedgehog. Endocrinology. 2010;151(3):1119–1128.

57. Ching S, Vilain E. Targeted disruption of Sonic Hedgehog in the mouse adrenal leads to adrenocortical hypoplasia. genesis. 2009;47(9):628–637.

58. Wood MA, et al. Fetal adrenal capsular cells serve as progenitor cells for steroidogenic and stromal adrenocortical cell lineages in *M. musculus*. Development. 2013;140(22):4522–4532.

59. Ahn S, Joyner AL. Dynamic Changes in the Response of Cells to Positive Hedgehog Signaling during Mouse Limb Patterning. Cell. 2004;118(4):505–516.

60. Gomez-Sanchez CE, et al. Development of monoclonal antibodies against human CYP11B1 and CYP11B2. Mol Cell Endocrinol. 2014;383(1–2):111–117.

61. Gancayco CA, et al. Intrinsic Adrenal TWIK-Related Acid-Sensitive TASK Channel Dysfunction Produces Spontaneous Calcium Oscillations Sufficient to Drive AngII (Angiotensin II)-Unresponsive Hyperaldosteronism. Hypertension. 2022;79(11):2552–2564.

62. Roybal K, et al. Mania-like behavior induced by disruption of CLOCK. Proceedings of the National Academy of Sciences. 2007;104(15):6406–6411.

63. Chen H, et al. PI3K-resistant GSK3 controls adiponectin formation and protects from metabolic syndrome. Proceedings of the National Academy of Sciences. 2016;113(20):5754–5759.

64. Taylor MJ, et al. Chemogenetic activation of adrenocortical Gq signaling causes hyperaldosteronism and disrupts functional zonation. Journal of Clinical Investigation. 2019;130(1):83– 93.

65. de Almeida Magalhaes T, et al. Extracellular carriers control lipid-dependent secretion, delivery, and activity of WNT morphogens. Dev Cell. 2024;59(2):244–261.e6.

66. Taipale J, et al. Effects of oncogenic mutations in Smoothened and Patched can be reversed by cyclopamine. Nature. 2000;406(6799):1005–1009.

67. Xu Q, et al. Vascular Development in the Retina and Inner Ear. Cell. 2004;116(6):883–895.

68. Westfall TA, et al. Wnt-5/pipetail functions in vertebrate axis formation as a negative regulator of Wnt/β-catenin activity. J Cell Biol. 2003;162(5):889–898.

69. Topol L, et al. Wnt-5a inhibits the canonical Wnt pathway by promoting GSK-3–independent β-catenin degradation. J Cell Biol. 2003;162(5):899–908.

70. Grumolato L, et al. Canonical and noncanonical Wnts use a common mechanism to activate completely unrelated coreceptors. Genes Dev. 2010;24(22):2517–2530.

71. Wu C, Nusse R. Ligand Receptor Interactions in the Wnt Signaling Pathway inDrosophila. Journal of Biological Chemistry. 2002;277(44):41762–41769.

72. Dong B, et al. Functional redundancy of Frizzled 3 and Frizzled 6 in planar cell polarity control of mouse hair follicles. Development. 2018;145(19):dev168468.

73. Wang Y, Guo N, Nathans J. The Role of Frizzled3 and Frizzled6 in Neural Tube Closure and in the Planar Polarity of Inner-Ear Sensory Hair Cells. The Journal of Neuroscience. 2006;26(8):2147–2156.

74. Dranow DB, Le Pabic P, Schilling TF. The non-canonical Wnt receptor Ror2 is required for cartilage cell polarity and morphogenesis of the craniofacial skeleton in zebrafish. Development. 2023;150(8). 10.1242/dev.201273.

75. Habas R, Kato Y, He X. Wnt/Frizzled Activation of Rho Regulates Vertebrate Gastrulation and Requires a Novel Formin Homology Protein Daam1. Cell. 2001;107(7):843–854.

76. Habas R, Dawid IB, He X. Coactivation of Rac and Rho by Wnt/Frizzled signaling is required for vertebrate gastrulation. Genes Dev. 2003;17(2):295–309.

77. Yamamoto S, et al. Cthrc1 Selectively Activates the Planar Cell Polarity Pathway of Wnt Signaling by Stabilizing the Wnt-Receptor Complex. Dev Cell. 2008;15(1):23–36.

78. Strutt DI, Weber U, Mlodzik M. The role of RhoA in tissue polarity and Frizzled signalling. Nature. 1997;387(6630):292–295.

79. Eubelen M, et al. A molecular mechanism for Wnt ligand-specific signaling. Science (1979). 2018;361(6403). 10.1126/science.aat1178.

80. Abu-Elmagd M, Garcia-Morales C, Wheeler GN. Frizzled7 mediates canonical Wnt signaling in neural crest induction. Dev Biol. 2006;298(1):285–298.

81. Martin E, Ouellette M-H, Jenna S. Rac1/RhoA antagonism defines cell-to-cell heterogeneity during epidermal morphogenesis in nematodes. Journal of Cell Biology. 2016;215(4):483–498.

82. Huang Y, Winklbauer R. Cell cortex regulation by the planar cell polarity protein Prickle1. Journal of Cell Biology. 2022;221(7). 10.1083/jcb.202008116.

83. Yang Y, Mlodzik M. Wnt-Frizzled/Planar Cell Polarity Signaling: Cellular Orientation by Facing the Wind (Wnt). Annu Rev Cell Dev Biol. 2015;31(1):623–646.

84. Wen J, et al. Loss of Dact1 Disrupts Planar Cell Polarity Signaling by Altering Dishevelled Activity and Leads to Posterior Malformation in Mice. Journal of Biological Chemistry. 2010;285(14):11023– 11030.

85. O’Connell AE, et al. Neonatal-Onset Chronic Diarrhea Caused by Homozygous Nonsense WNT2B Mutations. The American Journal of Human Genetics. 2018;103(1):131–137.

86. Zhang YJ, et al. Novel variants in the stem cell niche factor WNT2B define the disease phenotype as a congenital enteropathy with ocular dysgenesis. European Journal of Human Genetics. 2021;29(6):998–1007.

87. Berthon A, et al. WNT/β-catenin signalling is activated in aldosterone-producing adenomas and controls aldosterone production. Hum Mol Genet. 2014;23(4):889–905.

88. Valenta T, et al. Wnt Ligands Secreted by Subepithelial Mesenchymal Cells Are Essential for the Survival of Intestinal Stem Cells and Gut Homeostasis. Cell Rep. 2016;15(5):911–918.

89. Goss AM, et al. Wnt2/2b and beta-catenin signaling are necessary and sufficient to specify lung progenitors in the foregut. Dev Cell. 2009;17(2):290–8.

90. Farin HF, Van Es JH, Clevers H. Redundant sources of Wnt regulate intestinal stem cells and promote formation of Paneth cells. Gastroenterology. 2012;143(6):1518–1529.e7.

91. Cho S-H, Cepko CL. Wnt2b/beta-catenin-mediated canonical Wnt signaling determines the peripheral fates of the chick eye. Development. 2006;133(16):3167–77.

92. Kubo F, Takeichi M, Nakagawa S. Wnt2b controls retinal cell differentiation at the ciliary marginal zone. Development. 2003;130(3):587–98.

93. Lienkamp SS, et al. Vertebrate kidney tubules elongate using a planar cell polarity–dependent, rosette-based mechanism of convergent extension. Nat Genet. 2012;44(12):1382–1387.

94. Tarchini B, Lu X. New insights into regulation and function of planar polarity in the inner ear. Neurosci Lett. 2019;709:134373.

95. Pascoe L, et al. Mutations in the human CYP11B2 (aldosterone synthase) gene causing corticosterone methyloxidase II deficiency. Proceedings of the National Academy of Sciences. 1992;89(11):4996–5000.

96. Kranz A, et al. An improved Flp deleter mouse in C57Bl/6 based on Flpo recombinase. Genesis. 2010;48(8):512–520.

97. Turcu AF, et al. Comprehensive Analysis of Steroid Biomarkers for Guiding Primary Aldosteronism Subtyping. Hypertension. 2020;75(1):183–192.

98. Bray NL, et al. Near-optimal probabilistic RNA-seq quantification. Nat Biotechnol. 2016;34(5):525– 527.

99. Soneson C, Love MI, Robinson MD. Differential analyses for RNA-seq: transcript-level estimates improve gene-level inferences. F1000Res. 2016;4:1521.

100. Robinson MD, McCarthy DJ, Smyth GK. edgeR: a Bioconductor package for differential expression analysis of digital gene expression data. Bioinformatics. 2010;26(1):139–140.

101. Leek JT, et al. The sva package for removing batch effects and other unwanted variation in high-throughput experiments. Bioinformatics. 2012;28(6):882–883.

102. Ritchie ME, et al. limma powers differential expression analyses for RNA-sequencing and microarray studies. Nucleic Acids Res. 2015;43(7):e47–e47.

103. Yu G, et al. clusterProfiler: an R Package for Comparing Biological Themes Among Gene Clusters. OMICS. 2012;16(5):284–287.

104. Hao Y, et al. Dictionary learning for integrative, multimodal and scalable single-cell analysis. Nat Biotechnol. 2024;42(2):293–304.

105. Hippen AA, et al. miQC: An adaptive probabilistic framework for quality control of single-cell RNA-sequencing data. PLoS Comput Biol. 2021;17(8):e1009290.

106. McGinnis CS, Murrow LM, Gartner ZJ. DoubletFinder: Doublet Detection in Single-Cell RNA Sequencing Data Using Artificial Nearest Neighbors. Cell Syst. 2019;8(4):329–337.e4.

107. Linderman GC, et al. Zero-preserving imputation of single-cell RNA-seq data. Nat Commun. 2022;13(1):192.

108. Korsunsky I, et al. Fast, sensitive and accurate integration of single-cell data with Harmony. Nat Methods. 2019;16(12):1289–1296.

109. Traag VA, Waltman L, van Eck NJ. From Louvain to Leiden: guaranteeing well-connected communities. Sci Rep. 2019;9(1):5233.

110. Wierbowski BM, et al. Hedgehog Pathway Activation Requires Coreceptor-Catalyzed, Lipid-Dependent Relay of the Sonic Hedgehog Ligand. Dev Cell. 2020;55(4):450–467.e8.

111. Medina F, et al. Activated RhoA Is a Positive Feedback Regulator of the Lbc Family of Rho Guanine Nucleotide Exchange Factor Proteins. Journal of Biological Chemistry. 2013;288(16):11325– 11333.

